# Integrative Whole-Genome Analysis reveals Genomic Signatures of Innate Immunity in Indicine Cattle

**DOI:** 10.64898/2026.01.21.700751

**Authors:** Menaka Thambiraja, Mayank Roshan, Dheer Singh, Suneel Kumar Onteru, Ragothaman M. Yennamalli

## Abstract

Indicine cattle (*Bos indicus*) are known for resilience to infectious diseases and environmental stress. However, the genomic basis underlying this advantage remains poorly understood. To characterize variations in immune-related genetic elements in four indicine breeds (Kangayam, Gir, Tharparkar, and Sahiwal), a taurine breed (Holstein Friesian), and a taurine–indicine crossbreed (Karan Fries), we performed integrative whole-genome analysis. Using whole-genome data representing 108 animals, we identified structural variants, copy number variants, single-nucleotide variations, and insertions/deletions. High-impact single-nucleotide variations in the key innate immune genes, *CARD9* and *NLRP8*, shared across all indicine breeds, were absent in the other breeds. Genetic differentiation analysis identified several innate immune genes showing strong divergence between the indicine breeds and the taurine breed. Selective sweep detection analysis highlighted multiple breed-specific immune-related sweep regions. Functional enrichment analysis showed significant enrichment of immune pathways in indicine breeds. A comparison of the candidate genes with basal gene expression profiles of unchallenged peripheral blood mononuclear cells indicated that genomic variation influences the differential expression of several genes in indicine breeds. We synthesized the data from population genome structure analysis, nucleotide diversity, genetic differentiation, selective sweep analysis, and correlations with gene expression profiles. Indicine breeds exhibited a higher number of immune-related variants and stronger signals of selection in immune pathways. These findings provide a curated set of innate immune gene candidates in indicine breeds for future functional studies and breeding programs.

## Introduction

Cattle (Bos spp.) contribute substantially to dairy production, meat yield, draught power and rural livelihoods and are, therefore, economically significant worldwide. Cattle are broadly classified into two subspecies, *Bos taurus* (taurine/exotic) and *Bos indicus* (indicine), based on their origin, traits and adaptations. The distinct evolutionary histories, physiological characteristics, and genomic architectures of the two subspecies underpin differences in their productivity, adaptability, and immune competence. While taurine breeds, developed in temperate regions, have been selectively bred for productivity traits such as milk and meat [1], indicine cattle evolved under tropical climates, enabling them to tolerate heat, nutritional stress, and infectious diseases. Therefore, indicine breeds demonstrate greater thermotolerance, resilience to environmental pressures, and better disease resistance [2] than taurine breeds.

To combine the high milk productivity of taurine cattle with the resilience and hardiness of indicine breeds, crossbreeding programs have been undertaken. For example, Karan Fries of taurine-indicine lineage was developed by crossing Holstein Friesian, a taurine breed, with Tharparkar, an indicine breed [3]. However, despite improvement in production traits, the immune response and disease resistance of crossbred specimens do not consistently match those of pure indicine breeds.

The enhanced pathogen resistance and adaptability of indicine breeds are reported to lie in genomic elements related to unique immunity. Crossbreeding strategies to improve cattle health and productivity under tropical conditions must, therefore, consider the genomic basis of immunity. Successful strategies for the purpose require a detailed investigation of genomic signatures related to innate immunity, such as structural variants (SVs), copy number variants (CNVs) and single-nucleotide variants (SNVs) [4].

The mammalian immune system, including that of bovines, comprises innate and adaptive arms, each essential for host defense. The innate immune system, consisting of physical barriers, pattern-recognition receptors (PRRs), macrophages, neutrophils, and complement proteins, provides rapid, non-specific protection against invading pathogens [5]. In contrast, the adaptive immune system offers long-lasting, pathogen-specific responses driven by B and T cells, antigen presentation, somatic recombination and immunological memory. The innate and adaptive immune systems act synergistically to maintain immune homeostasis. Variation in genes associated with these immune components and pathways can significantly influence disease susceptibility and resilience in cattle. Understanding genomic diversity in immune genes is, therefore, crucial for identifying genetic markers, functional variants, and candidate genes that can enhance disease resistance through precision breeding.

Functionally important variants in coding and regulatory regions that modulate immune-related mechanisms. including host–pathogen interactions, antigen processing, cytokine signaling, and stress response mechanisms, can be identified by variant prediction based on whole genome sequencing (WGS). In cattle, studies use WGS to identify genes for immune traits and adaptive responses. In indicine cattle, genome-wide analyses highlight genes encoding toll-like receptors, as well as major histocompatibility, interferon, and chemokine receptors that have undergone natural selection for adaptation to tropical diseases [6]. Studies on admixed African cattle populations showed that candidate-selected genomic regions of indicine origin are associated with immune responses and heat tolerance, whereas a taurine origin is linked to inflammatory response. Selective signatures in immune-related genes, such as *MATR3, MZB1, STING1,* and *ATG4B*, have been associated with resistance to tick-borne diseases. *CARD11,* a key signaling molecule in both innate and adaptive immunity, has also been identified under selection [7]. In Zebu cattle, structural variant–focused WGS studies report duplications and deletions in genes associated with antimicrobial response, parasite resistance, and inflammatory signaling [8]. These findings demonstrate that high-resolution genomic data play a crucial role in unravelling evolutionary pressures that shape immune traits across cattle populations.

The comprehensive whole-genome variation profiling of multiple breeds can help identify candidate genes, adaptive variants, and genomic markers associated with immunity. Immune-related genes and allele-specific expression patterns distinguishing taurine and indicine cattle have been identified by previous studies that integrated genomic and transcriptomic data, using tissue-specific analyses of spleen and liver, with focus on single-nucleotide polymorphisms (SNPs) [9]. Our study extends this framework, using whole genome scale analysis across four indicine breeds, one crossbreed, and a taurine breed from geographically distinct locations, incorporating sequence-level variants as well as structural and copy number variants. We correlated these genomic variants with gene expression profiles in peripheral blood mononuclear cells (PBMCs) from healthy indicine breeds and crossbreeds, linking genomic variation to immune-related transcriptional differences. This integrative approach provides a broad view of the genomic architecture underlying immune adaptation and disease resistance in indicine cattle. Such insights directly support the development of climate-resilient, disease-tolerant, and high-performing cattle for breeding programs.

To explore the genetic diversity of immune-related genes in indicine cattle, we used whole-genome sequencing for the precise detection of SNVs, small insertions and deletions (InDel), SVs, CNVs, and selective sweep signatures across the entire genome. We focused on the comparative genomic analysis of indicine breeds, a crossbreed and a taurine breed, Holstein Friesian (HF), a dominant breed in industrial dairy farming worldwide. Out of the 50 officially recognized indicine breeds, we selected four common breeds: the dairy breeds, Gir (GR) and Sahiwal (SW), a draught cattle breed, Kangayam (KG), and a dual-purpose breed, Tharparkar (TP). A sixth breed, Karan Fries (KF), crossbred between indicine and exotic, was included for cross-reference.

Across the six breeds, using the variant callers, DELLY [10], GATK [11], and FreeBayes [12], we identified structural variants and small variants on the whole-genome pooled sequences of 108 samples. Population genomics analyses, using principal component analysis (PCA), the neighbor-joining tree [13], admixture, nucleotide diversity (π), the fixation index (F_ST_), absolute divergence (d_XY_), and selective sweep detection (raised accuracy in sweep detection, RAiSD) [14], revealed distinct genetic structures and strong signatures of selection linked to immune-related genes. Integrating these genome-wide analyses, we predicted potential innate immune markers. Correlating these markers with gene expression profiles of PBMCs under healthy condition revealed that, compared with KF, the indicine breeds, GR, TP, and SW, exhibited differential expression of several innate immune-related genes, indicating that these genes not only carry genomic variants but also display distinct basal immune responsiveness even under unchallenged conditions. Overall, our findings highlight the distinct genetic architecture of immune-related traits in indicine cattle, providing a foundation for identifying key immune markers to enhance disease resistance for sustainable breeding strategies.

## Materials and Methods

### Ethical statement

Ethical approval was obtained from the Institutional Animal Ethics Committee (IAEC) of the Indian Council of Agricultural Research (ICAR) - National Dairy Research Institute (NDRI), Karnal-132001, Haryana, India, for collecting blood and semen samples from the selected cattle breeds to isolate DNA for performing whole genome sequencing (Approval No. 46-IAEC-20-12).

### Sample collection and whole genome sequencing

We included six cattle breeds: four indicine breeds (KG, GR, TP, and SW), one crossbreed (KF), and one taurine breed (HF). Six unrelated healthy male animals from each breed were selected from three geographically distinct locations per breed across India, representing regional genetic diversity. Blood samples were collected directly from KG, GR, TP, SW and HF by venipuncture under standard veterinary procedures. For KF, both fresh and cryopreserved semen samples were used. Thus, we obtained 108 samples (6 individuals × 6 breeds × 3 locations = 108 samples) (Supplementary Figure 1). Samples collected from three different locations are denoted as L1 (Location 1), L2 (Location 2), and L3 (Location 3) for each breed.

Genomic DNA was isolated from the blood and semen samples and quantified using a Qubit assay kit. Genomic DNA samples from each breed were pooled based on location to prepare a breed-specific DNA set. Each pooled sample represented a single breed, comprising six animals from the same location, resulting in 18 pooled samples (1 pooled sample × 6 breeds × 3 locations). Supplementary Table 1 lists the source of the samples from each breed across the three different locations. The pooled samples were sequenced using the Illumina NovaSeq 6000 platform with 10X coverage. Experimental work, such as DNA isolation and quantification, was carried out at ICAR-NDRI. DNA sequencing was performed by RedCliffe Labs, Noida, India.

### Processing and mapping sequencing reads

Paired-end whole genome sequencing reads were obtained from RedCliffe Labs, Noida. Initial quality assessment was performed using FastQC [15], followed by the removal of adaptor sequences and low-quality bases using Trimmomatic in paired-end mode (TrimmomaticPE) [16] with settings as described below.

The *ILLUMINACLIP* parameter (Adapters.fasta:2:30:10:2:True) was used to identify and remove Illumina adapter sequences from the reads. In this setting, up to two mismatches were allowed when matching the adapter sequence. A palindrome clip threshold of 30 and a simple clip threshold of 10 were applied to ensure accurate detection of adapter contamination. Clipping was performed in forward and reverse orientations. The *LEADING:25* and *TRAILING:25* parameters were applied to remove low-quality bases from the beginning and end of each read, respectively. Bases with Phred scores below 25 were trimmed, ensuring that poor-quality regions at read extremities did not interfere with downstream analysis. Dynamic trimming was performed by the *SLIDINGWINDOW:4:15* option by scanning the read with a 4-base window and trimming the sequence once the average quality of bases in the window fell below a Phred score of 15. This helped remove regions where quality gradually declined. The *MINLEN:36* parameter ensured that only reads with a minimum length of 36 base pairs after trimming were retained for further analysis, preventing very short and potentially unreliable reads from being included. The *AVGQUAL:25* filter retained only reads with an average Phred quality score of at least 25, indicating overall high base-call accuracy across the entire read. *MAXINFO:36:0.5* was used to maximize the retention of information-rich bases while trimming. This parameter balanced read length and quality by targeting a minimum length of 36 bases with a strictness value of 0.5, helping preserve informative regions while discarding poor-quality segments.

The resulting high-quality reads were mapped to the *Bos taurus* reference genome (Hereford breed, ARS-UCD2.0) using BWA-MEM [17] with default settings. Alignment files were subsequently processed using SAMtools [18], including query-name sorting, mate-pair correction using samtools fixmate, coordinate sorting, indexing, duplicate marking using markdup, and final re-indexing to enable rapid access to specific genomic regions. Alignment and mapping quality (MAPQ) metrics were generated using samtools stats and samtools view. To ensure high-confidence alignments, we retained reads with MAPQ ≥ 20 (corresponding to a 99% probability of correct mapping) for variant calling.

### Variant calling with multiple tools

Genomic variants were identified using DELLY [10] for SVs and CNVs. DELLY detected large SVs such as translocations, inversions, insertions, deletions, duplications, and CNVs. FreeBayes [12] and GATK [11] were used for small variants such as SNVs and InDels. FreeBayes, a bayesian genetic variant detector, was used to call SNVs, InDels, multi-nucleotide variations (MNVs), and complex variants (combinations of multiple variant types) [12]. The GATK HaplotypeCaller identified high-confidence SNVs and InDels through local *de novo* reassembly at variant sites, without relying on existing mapping information [11].

All tools were used with default parameters. Variants were called by the selected tools across all genomic regions, including autosomes (29 chromosomes), X and Y chromosomes, the mitochondrial genome, and 1926 unplaced scaffolds, encompassing protein-coding genes, tRNAs, and other unannotated regions.

To remove low-quality variants, GATK variants were filtered using the following VariantFiltration expressions: SNVs failing QualByDepth (QD) < 2.0, FisherStrand (FS) > 60.0, RMSMappingQuality (MQ) < 40.0, StrandOddsRatio (SOR) > 3.0, MappingQualityRankSumTest (MQRankSum) < –12.5, and ReadPosRankSumTest (ReadPosRankSum) < –8.0.

Indels failing QD < 2.0, FS > 200.0, SOR > 10.0, and ReadPosRankSum < –20.0 were excluded (https://gatk.broadinstitute.org/hc/%20en-us/articles/360035531112%E2%80%93How-to-Filter-variants-either-with-%20VQSR-or-by-hard-filtering). FreeBayes variants were filtered to retain only those with QUAL ≥ 30 and MQ ≥ 20.

### Annotation using snpEff and the Ensembl Variant Effect Predictor

To ensure format consistency and accuracy, before annotation, all variants were normalized and validated using bcftools norm [18], VCF-validator, and VCF Assembly Checker. Using bcftools, SNVs for each breed were annotated with the latest *Bos taurus* reference SNP IDs (rsID) from Ensembl release 115. SnpEff [19] was used to annotate SVs and small variants into high-, moderate-, low-, and modifier-impact variants. High-impact variants included disruptive changes such as stop-gains, start-loss, exon deletions, or splice-site disruptions. Moderate-impact variants included missense variants and frameshift indels. Modifier variants are primarily located in non-coding or regulatory regions, while low-impact variants (such as synonymous variants) are generally neutral with no functional effect [20].

For downstream analysis, we focused on high-, moderate-, and modifier-impact variants with heterozygous (0/1) and homozygous (1/1) genotypes in genes, particularly immune-related genes. Genotype information was interpreted based on zygosity, the degree of similarity between alleles. The presence of a reference allele and an alternate allele are indicated by 0/1 while 1/1 indicates that both alleles differ from the reference, suggesting potential functional consequences due to complete alteration at that locus.

Since SnpEff does not support CNV annotation, CNVs were annotated using the Ensembl Variant Effect Predictor [21]. The CNVs were then categorized based on copy number (CN) into copy number gain (CPG; CN > 2) and copy number loss (CPL; CN < 2). Following annotation, immune-related genes were extracted using two complementary approaches: (1) annotation against the InnateDB database [22] to identify innate immune genes, and (2) keyword-based searches targeting immune-related terms to capture additional immune genes not listed in InnateDB by using an in-house Python script. In-house Python scripts were also used to identify common variants shared across the four indicine breeds, KG, GR, TP, and SW.

### Population genome structure and genetic diversity analysis

SNVs from the 18 pooled samples were merged into a unified dataset. High-confidence SNVs were filtered using Vcftools [23], retaining variants with SNV call rate > 90%, and minor allele frequency (MAF) < 0.05. Genetic differentiation among indicine, taurine, and crossbreed was evaluated by computing the pairwise Weir and Cockerham’s F_ST_ using Vcftools. A sliding-window scan with window size of 50 kb and a step size of 20 kb was used to detect genomic regions with genetic differentiation. A genome-wide F_ST_ analysis was performed using two complementary approaches. In the first approach, samples from the four indicine breeds (KG, TP, SW, and GR) were grouped and compared against the taurine breed (HF) and the crossbreed (KF) groups to study genetic differentiation among the three cattle groups. In the second approach, each indicine breed was individually compared with the taurine and the crossbreed groups (i.e., KG vs. HF vs. KF; GR vs. HF vs. KF; TP vs. HF vs. KF; and SW vs. HF vs. KF) to identify breed-specific patterns of genomic divergence. The top 1% of windows across autosomes showing the highest F_ST_ values were considered candidate selection regions and annotated using bedtools intersect [24] with *Bos taurus* (ARS-UCD2.0) gene coordinates. Within-breed genetic diversity was assessed using Vcftools, keeping the window and step size of nucleotide diversity (π) the same. All results were visualized using ggplot2 in R. In addition to F_ST,_ which measures relative genetic differentiation between populations, we evaluated absolute genetic divergence using d_XY_ statistics. Pairwise d_XY_ values were calculated using Pixy v2.0.0 [25], which compares nucleotide differences between two populations and accounts for both missing data and unequal sample sizes. A d_XY_ analysis was performed using a sliding window of 50kb, consistent with the window size used for F_ST_. Pairwise comparisons were carried out across each indicine breed with the taurine breed and the crossbreed separately. For each window, Pixy reported the average d_XY_ score along with the number of variant sites used in the calculation. Windows with zero informative sites were excluded from further analysis. To identify genomic regions showing both relative and absolute divergence, windows belonging to the top 1% of genomic-wide F_ST_ values were compared with the corresponding d_XY_ results. Regions showing high values in both analyses were considered breed-specific candidates.

The population genetic structure of the SNV genotypes identified across the six breeds was explored using PCA [26], admixture modelling [27], and phylogenetic tree construction. PCA was employed to reduce the dimensionality of the genetic data and visualize the distribution of genetic variation within and between the populations. Admixture analysis was performed to examine the genetic composition of the populations by estimating the proportion of ancestry derived from different source populations, providing insights into the historical admixture and shared ancestry among the six breeds.

The VCF file was converted to PLINK format (bed, bim, and fam) [28]. The variants were filtered using the built-in function in PLINK, with a MAF < 0.05 and missing genotype call rate < 10%. PCA was performed in PLINK by calculating eigenvectors across breeds and the first two principal components (PC1 vs. PC2) were visualized using ggplot2 in R. Admixture analysis was estimated using Admixture v1.3.0 across K = 1-7 and the optimal K value was selected based on the lowest cross-validation error, with the best predictive accuracy and an admixture plot was generated in R. A pairwise genetic-distance matrix was computed in PLINK, and used for constructing a neighbour-joining (NJ) phylogenetic tree, which was visualized in iTOL [29].

### Selective sweep detection

Selective sweep signals were detected using the µ statistic implemented in RAiSD [14], which integrates multiple signals of selection by jointly evaluating deviations in the site frequency spectrum, patterns of linkage disequilibrium, and reductions in genetic diversity across the genome. Genomic regions with strong selection signals within ±50 kb from each focal site were identified. These regions were annotated using bedtools with the *Bos taurus* reference genome (ARS-UCD2.0). Genes falling within the top 1% of the µ statistic distribution were considered potential candidates for selective sweeps. Candidate genes were subsequently subjected to KEGG pathway enrichment and gene ontology (GO) biological process analysis with the threshold of p-value < 0.05 using DAVID [30], [31].

### Correlation of genomic variants with gene expression profiles

Genomic variants, including SVs and CNVs from DELLY, small variants from FreeBayes and GATK, and candidate genes located within the top 1% F_ST_, d_XY_, and selective sweep regions identified by RAiSD, were integrated to generate a comprehensive list of putative innate immune genes specific to indicine breeds. These genes were compared with basal gene expression profiles derived from a previously published PBMC transcriptome dataset [32]. The referenced study generated transcriptome profiles of PBMCs collected from blood samples of three indicine breeds (GR, TP, and SW) and one crossbreed (KF), with three biological replicates per breed (n=3 per breed; total 12 animals). The sampled animals were examined by a qualified veterinarian to select clinically healthy specimens aged 12–16 months with no history of disease or infection. Blood samples were collected from the jugular vein under standardized conditions, and PBMCs were isolated using density gradient centrifugation. Total RNA extracted from the PBMCs was used for RNA-Seq library preparation and high-throughput sequencing. RNA-Seq data were processed following standard workflows, including quality assessment with FASTQC [15], adapter and low-quality base trimming with AdapterRemoval [33], mapping to the *Bos taurus* reference genome (ARS-UCD1.2) using HISAT2 [34], gene-level expression quantification with FeatureCounts [35], and differential expression analysis using edgeR [36] by comparing indicine breeds against the crossbreed group under unchallenged conditions. Innate immune genes identified from genomic variant and genetic differentiation analysis were correlated with differentially expressed genes to assess whether sequence-level variations were associated with transcriptional activity in PBMCs under unchallenged conditions. This integrative approach enabled the identification of genes showing both genetic divergence and functional expression differences, facilitating the detection of putative innate immune marker genes specific to indicine cattle.

## Results and Discussion

### Quality of sequence reads and alignment statistics

After trimming with Trimmomatic, all samples showed Q20 > 90% and Q30 > 85%. Therefore, only 3% of reads were discarded during filtering, indicating overall good sequencing quality. The summary of the statistics resulting from the whole genome sequence data before and after trimming is provided in Supplementary Table 2. Alignment statistics revealed that all the samples had more than 90% properly paired reads except GRL3 (89%). MAPQ was > 40 across all samples, while GRL3 showed a lower score (26.34). Since MAPQ > 20 corresponds to ∼99% confidence in correct alignment, all samples were considered suitable for variant analysis. Detailed metrics are provided in Supplementary Figure 2.

### Population structure and ancestry differentiation among indicine, taurine, and crossbreed cattle

PCA (Figure 1A) revealed a distinct clustering of indicine (KG, GR, TP, SW), taurine (HF), and crossbreed (KF) groups, with most indicine samples tightly grouped, except GRL3 and SWL2, which were slightly separated. Within the crossbreed, KF, the samples, KFL1 and KFL2, were closely clustered, with KFL3 slightly separated. HF and KF formed independent clusters, reflecting their exotic and mixed origins. However, HFL2 showed partial overlap with GRL3, the indicine sample, suggesting the possibility of limited introgression, or shared ancestry alleles.

**Figure 1.**
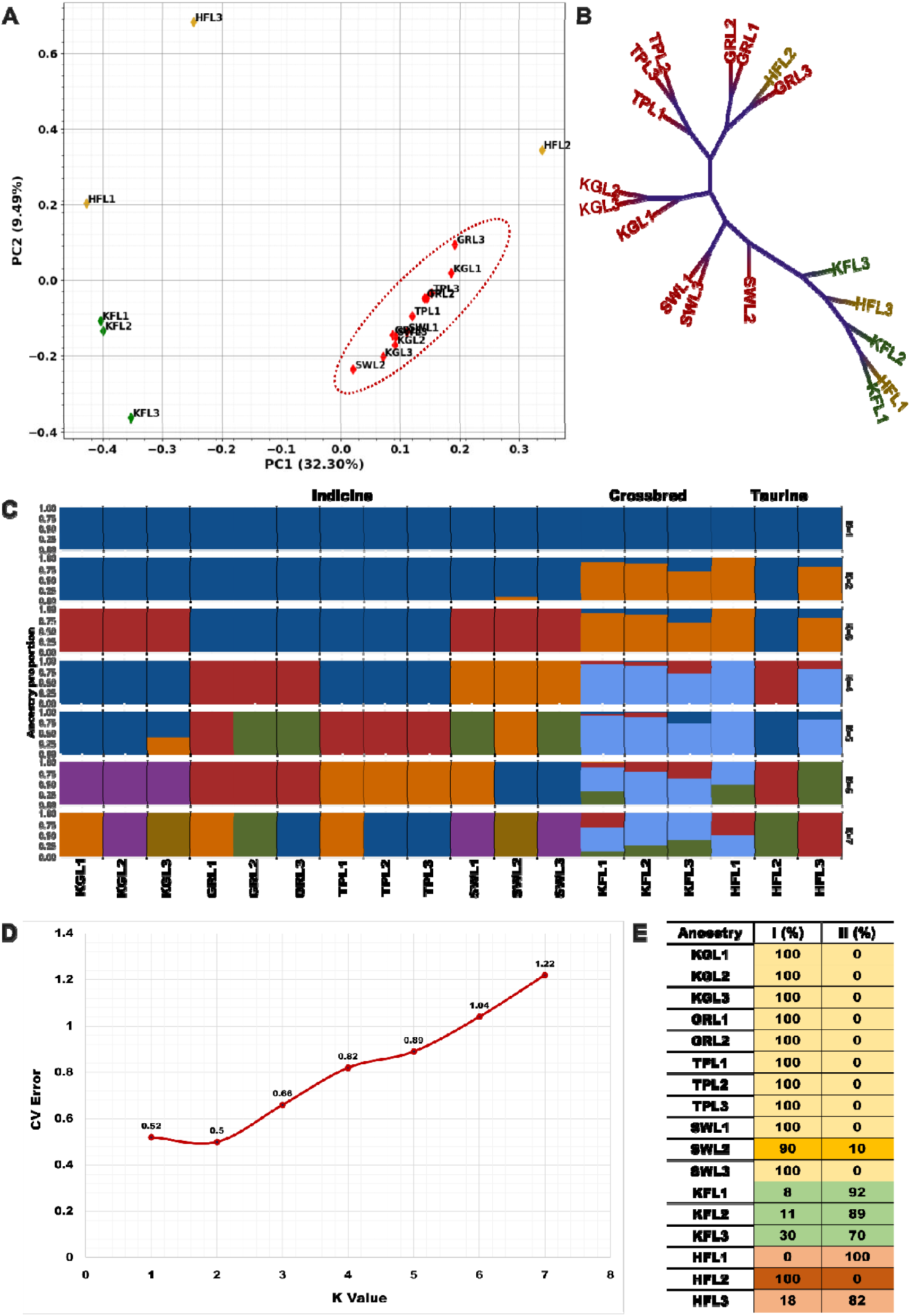
Population genome structure and admixture pattern across the three cattle groups. (A) Principal component analysis showing genetic clustering of the 18 pooled samples, with indicine breeds forming a distinct cluster (red dotted circle). Indicine (KG: Kangayam, GR: Gir, TP: Tharparkar, SW: Sahiwal), taurine (HF: Holstein Friesian), and crossbreed (KF: Karan Fries) samples are depicted as red, yellow, and green diamond markers, respectively. The three different locations from which the samples were taken are denoted as L1 (Location 1), L2 (Location 2), and L3 (Location 3). (B) Neighbour-joining phylogenetic tree illustrating genetic relationships among pooled samples. Indicine breeds (KGL1-KGL3, TPL1-TPL3, GRL1-GRL3, and SWL1-SWL3) are highlighted in red, crossbreeds (KFL1-KFL3) in green, and taurine (HFL1-HFL3) in yellow. (C) Admixture analysis highlighting genomic differentiation among indicine, taurine, and crossbreed groups. (D) Cross-validation error plot for admixture analysis of 18 pooled samples based on SNVs, with K values ranging from 1 to 7. (E) Ancestry proportions at K=2 across all 18 pooled samples, showing indicine (yellow), crossbreed (green), and taurine (red) ancestry components.

This pattern is supported by the NJ tree (Figure 1B), which reveals distinct clusters of each indicine breed, demarcating them from taurine, and crossbreed lineages. SWL2 has a slight overlap with taurine lineages, suggesting limited introgression. KF (TP × HF) was clustered closer with HF than with TP, indicating stronger taurine genetic contribution, consistent with earlier reports of taurine-dominant ancestry in crossbred cattle due to repeated backcrossing and selection for production traits [37, 38]. HFL2 was closely clustered with GRL3, reflecting a shared genetic background, limited introgression, or a technical error. Thus, PCA, as well as the NJ tree analysis based on whole-genome SNVs, highlights strong divergences between indicine and taurine breeds, a clear separation of population structure among the six cattle breeds and the three cattle groups with evidence of localized admixture.

Admixture analysis (Figure 1C) based on whole-genome SNVs clarified population structure and admixture pattern across 18 pooled samples. The cross-validation curve showed the lowest error at K = 2 (Figure 1D), indicating that the genomic variation can be best explained by two major ancestral groups: indicine and taurine lineages [39]. At K = 2 (Figure 1E), all four indicine breeds (KG, GR, TP, and SW) displayed nearly pure indicine ancestry, except SWL2, which showed 10% taurine admixture, indicating limited introgression. KF, a known TP × HF crossbreed, showed 70-92% taurine ancestry, reflecting its strong taurine genetic background, matching the NJ tree pattern clustering. Though HFL1 displayed 100% taurine ancestry and HFL3 displayed 82% taurine ancestry, HFL2 unexpectedly showed complete clustering with indicine ancestry, with no detectable taurine genetic component. Given the well-established genetic distinction between taurine and indicine cattle, this pattern is unlikely to reflect true biological admixture. Taken together, the consistent alignment of HFL2 with indicine samples across PCA, NJ tree, and admixture analyses strongly indicates a sample misidentification or labelling error during collection or processing. Hence, HFL2 was excluded and we performed subsequent analyses with 17 pooled samples. Overall, the admixture results confirm clear genetic differentiation between indicine and taurine lineages while highlighting admixture patterns in crossbreeds such as KF, reflecting both historical breeding strategies and genetic introgression events.

### Indicine-specific immune CNVs relative to taurine and crossbreed cattle

DELLY was used to detect CNVs, classified as copy number gain (CPG; CN > 2) and copy number loss (CPL; CN < 2). Across all breeds, CPL events were more frequent than CPG events (Figure 2A). CPLs (deletions) are reported to be shorter but occur more frequently, whereas CPGs (duplications) tend to span larger genomic regions but are less frequent [40]. CNV-associated genes were identified in pooled samples, and common CNV genes were determined within each breed.

**Figure 2.**
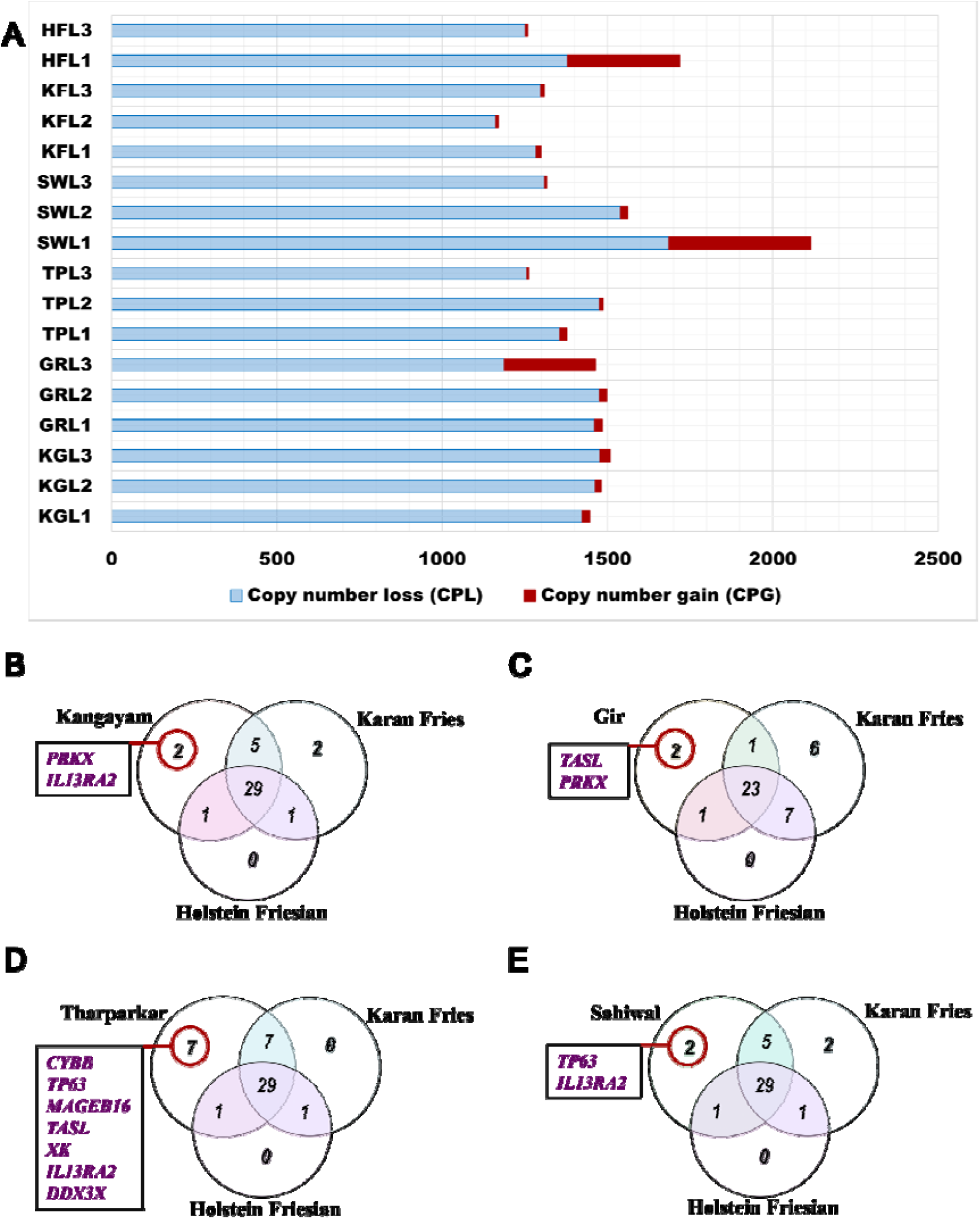
Copy number variants identified by DELLY. (A) Distribution of copy number variants across all 17 pooled samples, showing copy number loss and copy number gain. (B-E) Number of innate immune genes specific to Kangayam (B), Gir (C), Tharparkar (D), and Sahiwal (E) in comparison with Karan Fries and Holstein Friesian.

A comparative analysis of indicine, exotic, and crossbreed groups (Figure 2B-2E) revealed several innate immune genes uniquely affected by CNVs in indicine breeds. In KG (Figure 2B), CPLs in *IL13RA2* (interleukin 13 receptor alpha 2) and *PRKX* (protein kinase, X-linked) could suggest a potential impact on macrophage activation, antibody differentiation and myeloid cell (macrophage and granulocyte) development. In GR (Figure 2C), CPLs were identified in *TASL,* an adaptor present in toll-like receptors involved in antiviral immunity and *PRKX*. In TP (Figure 2E), CPLs were identified in multiple immune-related genes, including *CYBB* (involved in phagocytosis)*, TP63* (epithelial barrier integrity, antiviral defence)*, MAGEB6* (melanoma antigen family B16 member that stimulates immune response)*, TASL, XK, IL13RA2,* and *DDX3X* (antiviral defense and type-I interferon production). Similarly, in SW (Figure 2D), CPLs were observed in *TP63* and *IL13RA2*.

As KF is crossbred between TP and HF, several genes with CPLs were shared by both parental breeds. These included genes, shared with TP, involved in antigen presentation and T-cell activation through the major histocompatibility complex class-II (*BOLA-DQA5,* and *BOLA-DQB),* in the positive regulation of innate immune and antiviral response *(MID2,* also known as *TRIM1*), in immune response signaling (*VSIG1,* a member of the immunoglobulin superfamily), in autophagy (*ATG4A),* in interleukin-1 receptor family function (*IL1RAPL2),* and in the regulation of inflammatory response (*TSC22D3)*. CPLs in 30 immune genes were common to KF and HF. This pattern highlights the inheritance of CNV patterns from both parental lineages.

### Structural variants in immune-related genes are higher in indicine breeds

Across the 17 pooled samples, DELLY identified five classes of structural variants: duplications, deletions, inversions, insertions, and translocations (Figure 3A). Deletions and translocations were the most frequent, followed by inversions, duplications, and insertions. GRL3 had the highest number of translocations, and KFL2 had more inversions. To identify breed-specific variants, only genes with variants shared across all three locations within each breed were considered, resulting in six breed-specific gene sets (Figure 3B). The highest number of genes with duplications was found in TP (592 genes), followed by KG (425 genes). GR had a higher number of deletions (409 genes), and KF had a higher number of inversions (25 genes). HF had comparatively fewer genes with variants. Although translocations occurred frequently, only a few genes, all classified as high-impact, were annotated. The annotated genes with variants categorized by high-, moderate-, low-, and modifier-impact for all 17 samples are provided in Supplementary Table 3.

**Figure 3.**
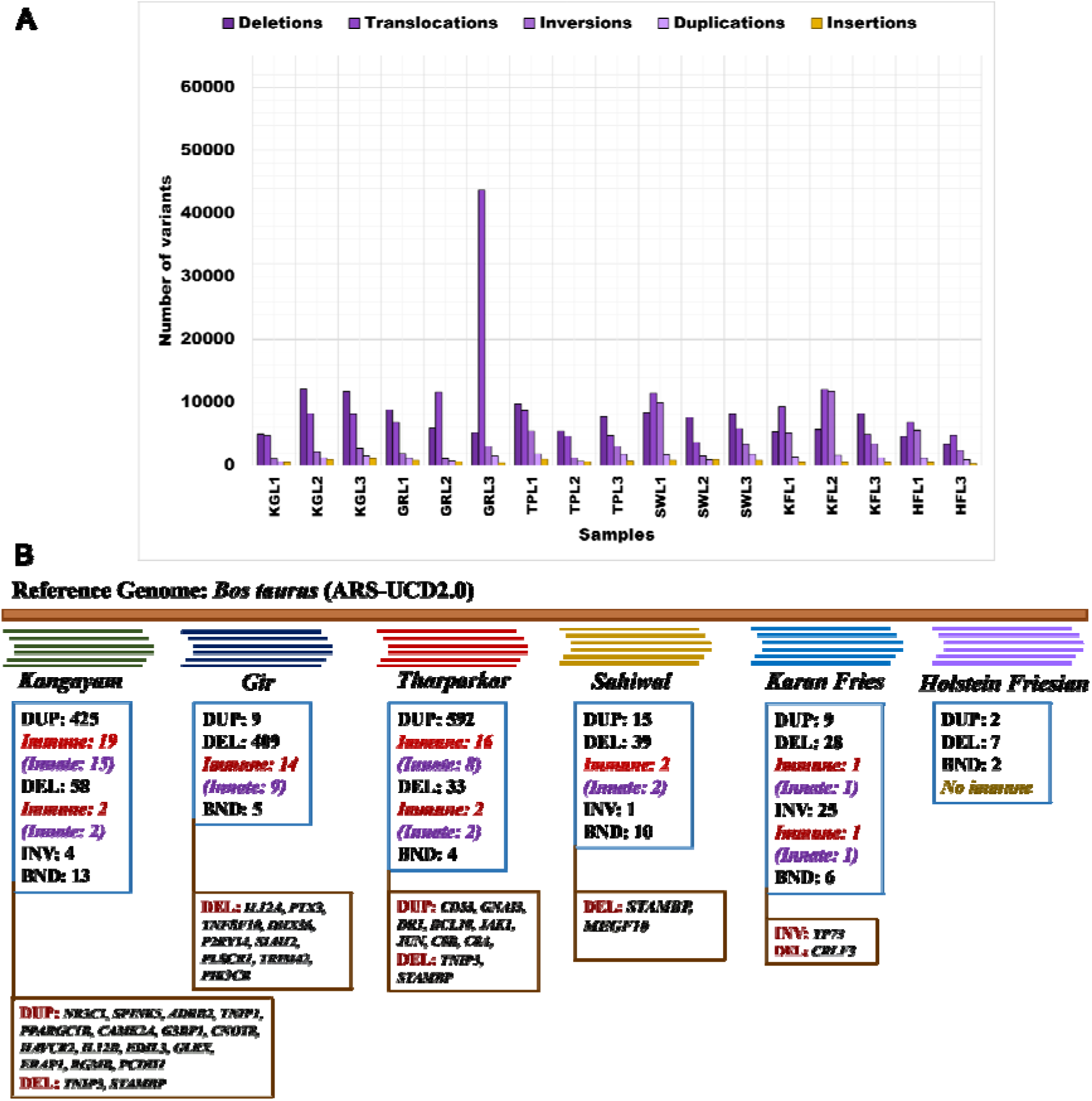
Structural variants (SVs) identified by DELLY. (A) Number of variants identified across all 17 pooled samples. (B) Number of genes, including innate immune genes (identified via InnateDB) and other immune-related genes (identified via keyword-based search) associated with these variants across all six breeds. Structural variants include DEL (deletion), DUP (duplication), INV (inversion), and BND (translocation).

SnpEff classified the genes with variants into high-, moderate-, modifier-, and low-impact categories (Table 1). A high number of high-impact variants, primarily translocations and deletions, were observed in GR, followed by KG, SW, KF, TP, and HF. Moderate-impact variants were mainly duplications in TP (588 genes), KG (417 genes), and inversions in KF (25 genes). Modifier-impact variants, typically located in regulatory and non-coding regions, were more common in indicine breeds (GR, KG, TP, SW) than in KF and HF.

**Table 1:** Number of genes with variants showing high-, moderate-, modifier-, and low-impact common across all three locations within each breed, identified by DELLY and annotated with SnpEff. The variants include DELs (deletions), DUPs (duplications), INVs (inversions), and BNDs (translocations) across the six breeds.

| Breed | High | Moderate | Modifier | Low |
| --- | --- | --- | --- | --- |
| Kangayam | DEL: 31 (*immune: 1*),<br>DUP: 2, BND: 12 | DEL: 5, DUP: 417<br>(*immune: 20*), INV: 4 | DEL: 50,<br>DUP: 56 | DEL: 6,<br>DUP: 1 |
| Gir | DEL: 408 (*immune: 14*),<br>DUP: 1, BND: 6 | DUP: 5 | DEL: 73,<br>DUP: 9 | DEL: 3 |
| Tharparkar | DEL: 18 (*immune: 1*),<br>DUP: 1, BND: 5 | DUP: 588 (*immune: 16*) | DEL: 25,<br>DUP: 101 | DEL: 1 |
| Sahiwal | DEL: 29, DUP: 2, BND: 1 | DEL: 2, DUP: 8, INV: 1 | DEL: 22,<br>DUP: 15 | DEL: 4,<br>DUP: 1 |
| Karan Fries | DEL: 20 (*immune: 1*),<br>DUP: 1, INV: 2, BND: 6 | DUP: 5, INV: 25<br>(*immune: 1*) | DUP: 9 | - |
| Holstein Friesian | DEL: 2, DUP: 2, BND: 2 | - | DEL: 7, DUP: 2 | DUP: 1 |

Breed-specific variant analysis revealed that indicine breeds had more genes with structural variants than KF and HF. Several of these variants were observed in immune-related genes across breeds (Figure 3B). KG had more immune genes with SVs, followed by TP and GR. In contrast, no immune-related variants shared across all three locations were detected in HF, highlighting reduced structural variability in immune genes.

In KG, duplications were identified in 15 innate immune genes with moderate-impact heterozygous variants (0/1). These genes are involved in the regulation of immune homeostasis (*NR3C1* and *PPARGC1B*), innate antiviral defense (*G3BP1*), the modulation of inflammatory responses (*CAMK2A, HAVCR2, CNOT8, EDIL3,* and *RGMB*), anti-inflammatory and antimicrobial protection (*SPINK5*), the negative regulation of inflammation (*TNIP1*), the activation of innate and adaptive immunity (*IL12B*), redox regulation and immune signaling (*GLRX*), cell-cell adhesion and barrier integrity (*PCDH1*) and the maintenance of immune tolerance (*ADRB2*).

*ERAP1* contributes to antigen processing during bacterial infection in cattle. Reduced expression of this gene showing protective responses to bovine tuberculosis, together with increased non-classical MHC class I molecule HLA-E expression and IFN-γ production, suggests that it is a potential biomarker signature for bTB resistance [41] (Figure 3B).

In Tharparkar (TP), we identified the duplication of eight genes (0/1) involved in key immune functions: complement activation (*C8A* and *C8B)*, regulation of T cells, NK cells, and induction of interferon production (*CD53*), autophagy signaling and pathogen clearance (*GNAI3*), IL-2–mediated signal transduction (*JAK1*), essential mediators of NF-kappaB activation and apoptosis induction (*BCL10*), negative regulation of antiviral innate responses (*DR1*), and regulation of immune cell activation and differentiation, particularly in T cells (*JUN*) (Figure 3B).

Both KG and TP had high-impact deletions with frameshift and splice-site heterozygous variants (0/1) in *TNIP3*, an innate immune gene involved in the negative regulation of NF-κB2 signaling and maintaining immune homeostasis [42]. In SW, deletions were observed in two immune genes (0/1), *STAMBP* and *MEGF10. MEGF10* is involved in the phagocytic clearance of apoptotic cells, an essential innate immune process that limits inflammation and maintains tissue homeostasis [43]. Frameshift deletions in *STAMBP* were observed in KG, TP, and SW samples, indicating a shared disruptive variant across these breeds. *STAMBP* encodes for a deubiquitinase (DUB) that acts as a central regulator of innate immunity. It negatively regulates the NLRP3 inflammasome and IL-1β secretion, preventing excessive inflammatory responses [44].

In GR, nine innate immune genes, with high-impact deletion (0/1), predicted to cause transcript ablation, were identified. These genes were related to the activation of macrophages, NK cells, and T cells (*IL12A)*, pathogen recognition *(PTX3)*, induction of apoptosis (*TNFSF10)*, sensing of viral component (*DHX36)*, the recruitment of macrophages and neutrophils, and the modulation of inflammatory response *(P2RY14)*, the regulation of T-regulatory cells (*SIAH2)*, the induction of type-I interferon (*PLSCR1)*, innate antiviral defense (*TRIM42)*, and signaling required for neutrophil activation and migration at the site of infection (*PIK3CB*) (Figure 3B).

In KF, a moderate-impact inversion was identified in *TP73* (1/1). This gene is involved in macrophage-mediated innate immunity, playing a complex role in innate and adaptive immunity and regulating inflammatory response. High-impact frameshift deletions were observed in *CRLF3* (cytokine receptor-like factor 3), an evolutionarily conserved gene in vertebrates that functions as a negative regulator of antiviral immune response [45].

Compared with other breeds, KG and TP exhibited a higher number of gene duplications. This finding is consistent with previous array comparative genomic hybridization analyses across indicine (KG, TP, GR, and SW), taurine (HF), and crossbred (KF) cattle that reported a greater number of duplicated innate immune genes in KG and TP. In contrast, KF and HF showed fewer genes with duplications and no deletions. These patterns suggest reduced genetic variability in the immune gene repertoire of KF and HF compared with indicine breeds [46].

### Small variant identification using FreeBayes and GATK with default and ploidy 12 parameters

Small variants were identified using two complementary variant-calling tools, FreeBayes and GATK. Cross-validation between the two tools helped improve confidence in the identified variants. FreeBayes and GATK were run with both default diploid settings and a ploidy value of 12 to account for the pooled samples (six animals per pool, totaling 12 alleles). The ploidy-12 approach enables the estimation of the number of individuals in a pool carrying a given variant, although its utility is constrained by the limited availability of downstream population-genomics tools that support high-ploidy data.

Before and after filtering, under the default (Figure 4A) as well as the ploidy-12 approach (Figure 4B), FreeBayes detected a higher number of variants than GATK. FreeBayes captured a broader spectrum of variant types, including SNVs, insertions, deletions, MNPs, and complex variants (combinations of multiple variant types). GATK, on the other hand, primarily identifies SNVs and InDels. Variants called by both tools were distributed across all genomic regions, including autosomes, X chromosomes, unplaced scaffolds, and the mitochondrial genome, spanning coding genes, tRNAs, and uncharacterized loci.

**Figure 4.**
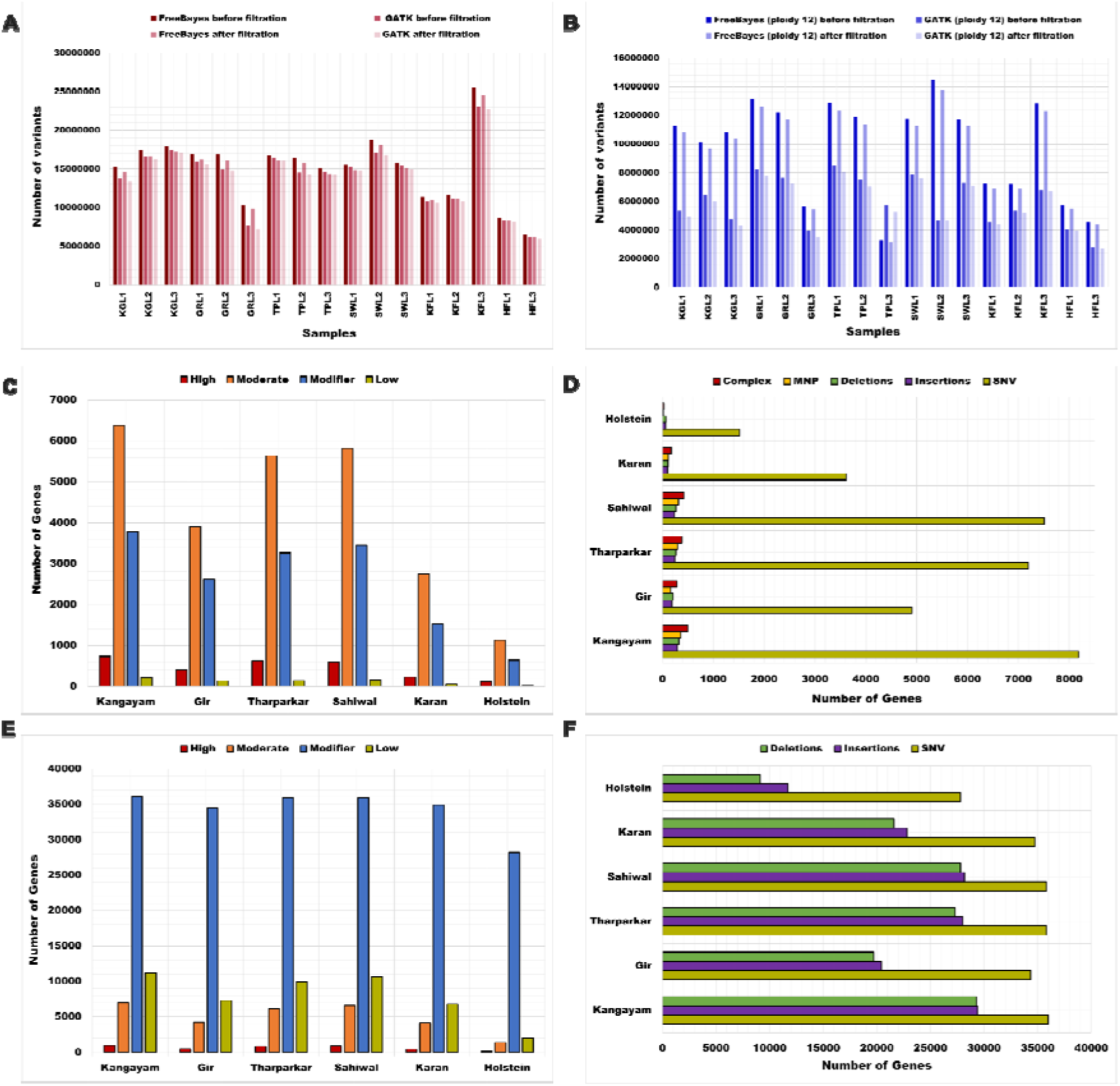
Total number of variants, genes carrying different types of variants, and their impacts identified by FreeBayes and GATK. A) Number of variants identified by FreeBayes and GATK with default parameters before and after applying filters. B) Number of variants identified by FreeBayes and GATK with ploidy12 before and after applying filters. (C) Number of genes carrying variants identified by FreeBayes with high-, moderate-, modifier-, and low-impacts. (D) Number of genes with different types of variants, such as SNVs, MNVs, insertions, deletions, and complex variants, identified by FreeBayes. (E) Number of genes carrying variants identified by GATK with high-, moderate-, -modifier, and low-impacts. (F) Number of genes with different types of variants, such as SNV, insertion, and deletion variants, identified by GATK.

As with DELLY, in FreeBayes and in GATK, breed-specific variants were identified by considering only those variants consistently present across all three locations within each breed. Although FreeBayes identified more variants than GATK, annotation with snpEff revealed that GATK-identified variants were annotated with a larger number of genes. We note that FreeBayes identified more moderate-impact variants (Figure 4C), reflecting the predominance of missense variants and in-frame InDels. In contrast, GATK detected a larger proportion of modifier-impact variants, indicating that many variants were located in non-coding and regulatory regions (Figure 4E). Both callers consistently identified SNVs as the most abundant variant type (Figures 4D and 4F). Both tools showed that KG had a higher number of variants, followed by SW, TP, and GR, while HF had fewer variants (Figures 4D and 4F). The annotated genes with high-, moderate-, modifier-, and low-impact variants for all 17 samples as identified by GATK are provided in Supplementary Table 4.

After quality filtration, 24% of FreeBayes variants and 2% of GATK variants were lost, suggesting that GATK inherently calls a greater proportion of high-quality variants. Therefore, variants with SNVs identified using GATK were mapped with the latest *Bos tauru*s reference SNP IDs (rsIDs), Figure 5.

**Figure 5.**
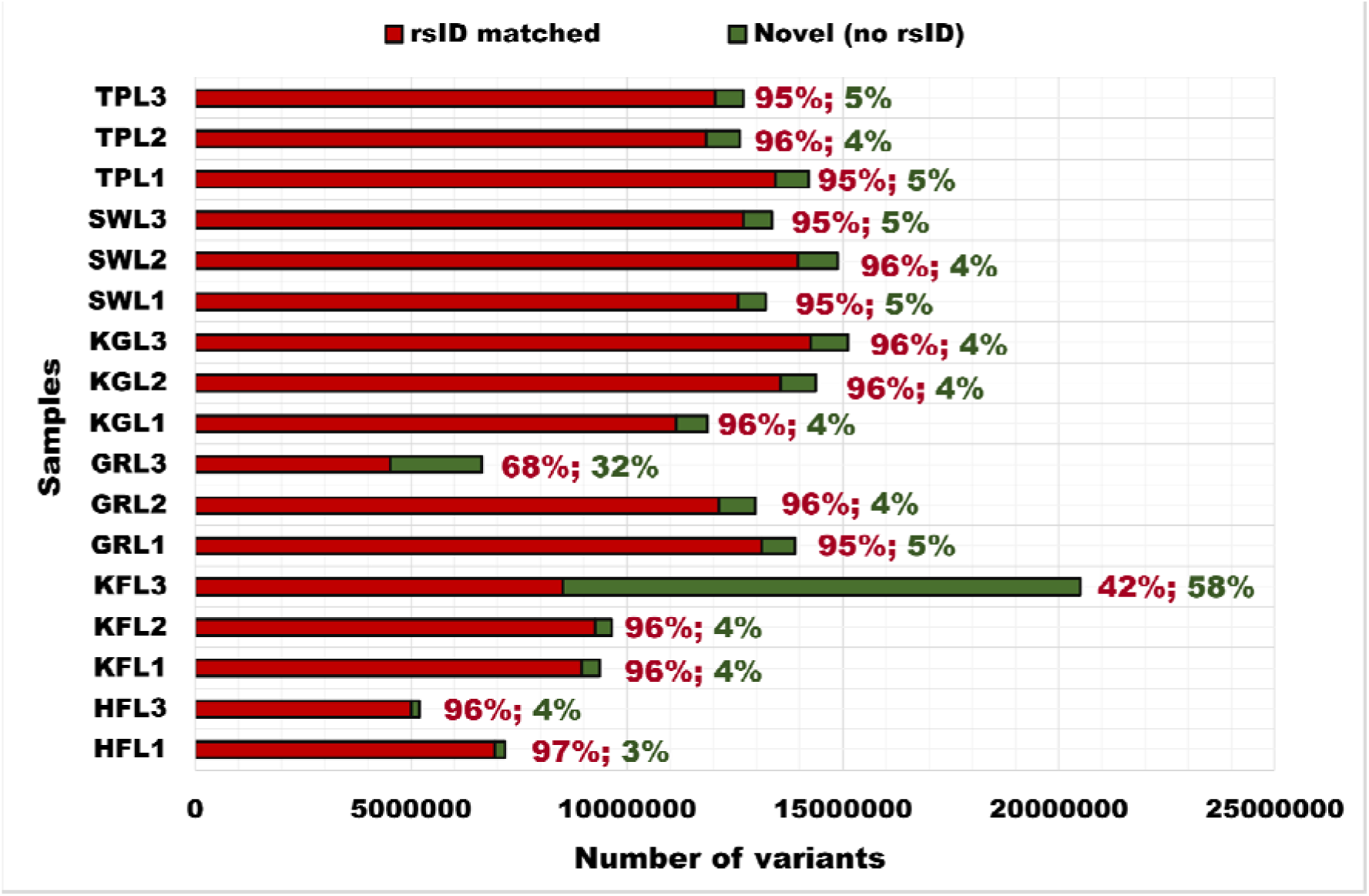
SNV variants identified by GATK matched with Bos taurus reference SNP ID (rsID). Samples of indicine (KG: Kangayam, GR: Gir, TP: Tharparkar, SW: Sahiwal), crossbreed (KF: Karan Fries), and exotic (HF: Holstein Friesian) breeds from three different locations (L1: Location 1, L2: Location 2, L3: Location 3) were aligned with the latest Bos taurus  reference SNP ID. The red bar represents the identified variants matched with rsID (Bos taurus variation), and the green bar represents the variants identified as unique (without rsID).

The resultant mapping revealed that 94–95% of variants across all indicine breeds matched existing rsIDs, where 4–5% were unique (with no match to rsIDs) except for GRL3, where only 68% of the variants matched with rsID and 32% were unique. KF, matched 96% of variants with rsIDs, except KFL3, where only 58% matched. The exotic breed, HFL1, showed 97% matches, and HFL3 showed 96% matches. Overall, variant calling by FreeBayes and GATK revealed that the indicine breeds, KG, GR, TP, and SW, had more variants than KF and HF when aligned with the *B. taurus* reference genome.

A higher number of genes were annotated with variants identified by GATK. Across both FreeBayes and GATK, the total number of genes with variants was consistently highest in KG, followed by SW, TP, GR, and KF, with HF showing the fewest (Table 2). For variants in innate immune genes, FreeBayes identified the most in TP and SW (233 genes each), followed by KG, GR, KF and HF, whereas GATK identified the highest number in SW (894 genes), followed by KG, TP, GR, KF, and HF. Both variant callers consistently indicated that indicine breeds harbored more variants in innate immune genes than the crossbreed and the taurine breeds. Among indicine breeds, SW, TP, and KG exhibited more variants in innate immune genes than GR, highlighting the importance of genomic differences in innate immune genes within indicine breeds.

**Table 2.**
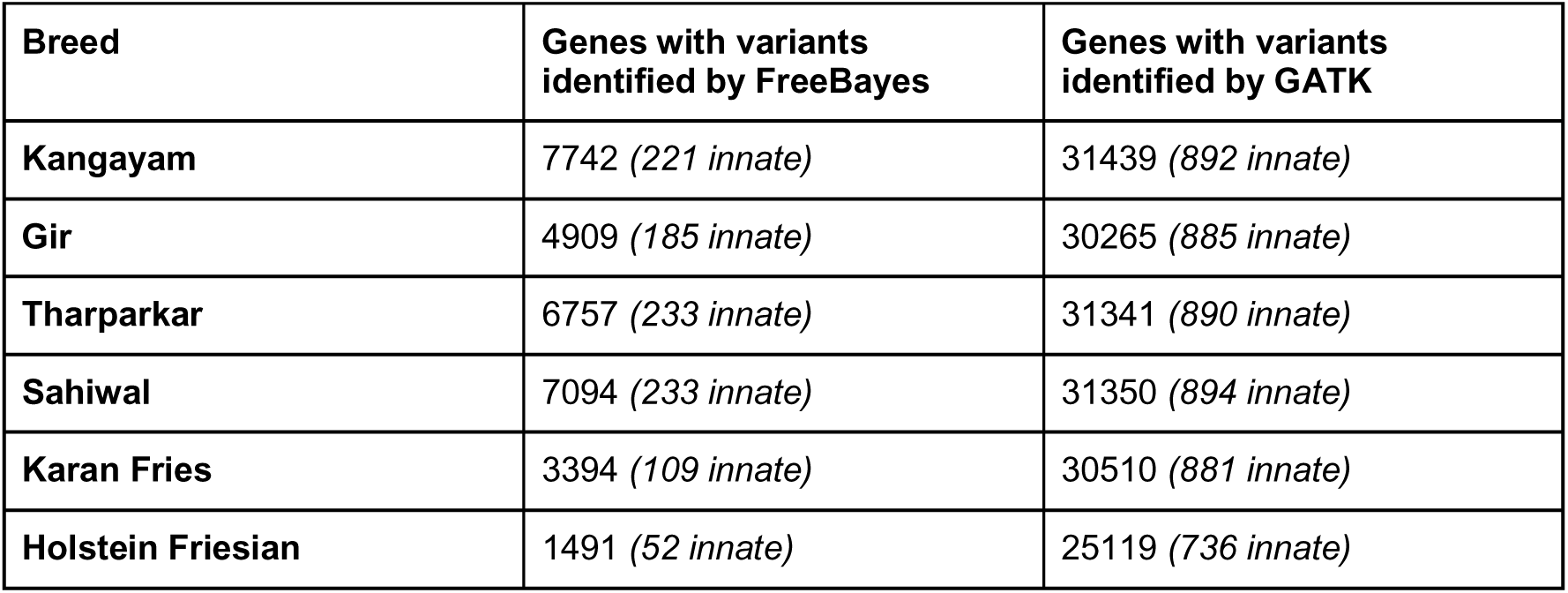
Number of genes and immune genes with variants identified by FreeBayes and GATK. Innate immune genes identified by variant callers are shown within parentheses and in italics.

| Breed | Genes with variants identified by FreeBayes | Genes with variants identified by GATK |
| --- | --- | --- |
| Kangayam | 7742 (*221 innate*) | 31439 (*892 innate*) |
| Gir | 4909 (*185 innate*) | 30265 (*885 innate*) |
| Tharparkar | 6757 (*233 innate*) | 31341 (*890 innate*) |
| Sahiwal | 7094 (*233 innate*) | 31350 (*894 innate*) |
| Karan Fries | 3394 (*109 innate*) | 30510 (*881 innate*) |
| Holstein Friesian | 1491 (*52 innate*) | 25119 (*736 innate*) |

Not all variants in immune-related genes have the same functional significance. Therefore, SnpEff was used to annotate the variants identified by FreeBayes and GATK, classifying them as high- (loss-of-function), moderate- (missense) and modifier- (regulatory and non-coding) impacts. Table 3 summarizes the moderate- and modifier-impact variants in immune genes, including innate immune genes extracted from InnateDB.

**Table 3.** Moderate- and modifier-impact immune genes identified by both FreeBayes and GATK across all breeds. The number of innate immune genes identified with the InnateDB gene list is shown within parentheses and in italics.

| Breed | Variants identified by FreeBayes | Variants identified by GATK | Variants identified by both FreeBayes and GATK |
| --- | --- | --- | --- |
| **Kangayam** | Moderate: 379<br>(204 *innate*)<br>Modifier: 200<br>(111 *innate*) | Moderate: 410<br>(225 *innate*)<br>Modifier: 1427<br>(892 *innate*) | Moderate: 327<br>(183 *innate*)<br>Modifier: 167<br>(95 *innate*) |
| **Gir** | Moderate: 267<br>(136 *innate*)<br>Modifier: 146<br>(87 *innate*) | Moderate: 296<br>(157 *innate*)<br>Modifier: 1371<br>(873 *innate*) | Moderate: 224<br>(118 *innate*)<br>Modifier: 127<br>(77 *innate*) |
| **Tharparkar** | Moderate: 350<br>(188 *innate*)<br>Modifier: 166<br>(97 *innate*) | Moderate: 389<br>(208 *innate*)<br>Modifier: 1418<br>(890 *innate*) | Moderate: 314<br>(169 *innate*)<br>Modifier: 148<br>(87 *innate*) |
| **Sahiwal** | Moderate: 357<br>(185 *innate*)<br>Modifier: 177<br>(95 *innate*) | Moderate: 392<br>(210 *innate*)<br>Modifier: 1422<br>(893 *innate*) | Moderate: 320<br>(171 *innate*)<br>Modifier: 153<br>(84 *innate*) |
| **Karan Fries** | Moderate: 170<br>(86 *innate*)<br>Modifier: 81<br>(40 *innate*) | Moderate: 239<br>(120 *innate*)<br>Modifier: 1361<br>(872 *innate*) | Moderate: 138<br>(70 *innate*)<br>Modifier: 63<br>(32 *innate*) |
| **Holstein Friesian** | Moderate: 67<br>(37 *innate*)<br>Modifier: 40<br>(21 *innate*) | Moderate: 90<br>(47 *innate*)<br>Modifier: 1019<br>(666 *innate*) | Moderate: 56<br>(34 *innate*)<br>Modifier: 28<br>(15 *innate*) |
| **All indicine (KG, GR, TP, and SW)** | Moderate: 73<br>(34 *innate*)<br>Modifier: 39<br>(21 *innate*) | Moderate: 73<br>(34 *innate*)<br>Modifier: 1115<br>(710 *innate*) | Moderate: 55<br>(24 *innate*)<br>Modifier: 32<br>(17 *innate*) |

Both callers consistently identified the highest number of moderate- and modifier-impact variants in innate immune genes in KG (183 and 167 innate genes), followed by SW (171 and 84 innate), TP (169 and 87 innate), GR (118 and 77 innate), and KF (70 and 32 innate), with HF showing the fewest variants (34 and 15 innate). These results show that indicine breeds (KG, SW, TP, and GR) exhibited more variants in immunity-related genes than KF and HF. The greater number of variants in immune genes in indicine breeds may reflect adaptation and enhanced disease resistance. Moderate- and modifier-impact variants in innate immune genes across indicine breeds are particularly important to study because they might affect immunity through distinct mechanisms. Moderate-impact variants, such as missense and in-frame changes, could directly influence protein function, whereas modifier-impact variants, primarily located in regulatory regions, are more likely to affect gene regulation and expression. Together, these variants may contribute to disease resistance and other economically relevant traits in indicine breeds.

### High-impact innate immune variants identified by both FreeBayes and GATK

In contrast to moderate- and modifier-impact variants, high-impact variants were less frequent in innate immune genes across indicine, taurine, and crossbreed cattle. However, similar to the pattern observed for moderate- and modifier-impact variants, high-impact variants were highly distributed in indicine breeds, whereas HF and KF had very few variants in innate immune genes (Table 4). FreeBayes identified a higher number of high-impact variants across innate immune genes in KG, followed by TP, GR, SW, KF, and HF. GATK identified more variants in TP, followed by SW, KG, KF, GR, and HF. To obtain high-confidence variant-associated genes, we focused on genes with variants identified by both callers. These resultant high-impact variants in innate immune genes were most frequent in TP (13 genes), followed by KG (11), SW and GR (9 each), KF (3), and HF (2). High-impact SNVs, insertions, and deletions identified by both variant callers occurred as either homozygous (1/1) or heterozygous (0/1) genotypes (Table 4).

**Table 4.** High-impact innate immune genes identified by FreeBayes and GATK across all breeds. Genes formatted in bold-italics represent those detected by both tools.

| Breed | Variants identified by FreeBayes | Variants identified by GATK |
| --- | --- | --- |
| Kangayam | SNV: *CXCR1*, *CCR6*, *CARD9*, *TRIM66*, *NLRP8*, *IGF1R* (1/1); *C5* (0/1)<br>INS: *TNIP1*, *PIAS1*, *IRF6*, *TBKBP1*, *TLR3*, *TOLLIP*, *XIAP* (1/1); *RXRA*, *TRIM66* (0/1)<br>DEL: *SPINK5*, *JAM3*, *ZFPM2* (1/1)<br>Complex: *HDAC1* (1/1) | SNV: *CXCR1*, *CCR6*, *CARD9*, *TRIM66*, *CD46*, *NLRP8*, *IGF1R*, *HDAC11* (1/1); *CD100LF*, *C1R*, *C5* (0/1)<br>INS: *IRF6*, *TOLLIP*, *NUMBL* (1/1)<br>DEL: *SPINK5*, *JAM3*, *GSK3A* (1/1) |
| Gir | SNV: *PIK3CB*, *PLG*, *CCR6*, *CARD9*, *NLRP8*, *MAPK8*, *SH2DIA* (1/1); *IRAK2*, *DEFB7* (0/1)<br>DEL: *SPINK5*, *ZFPM2*, *JAM3*, *GSK3A* (1/1) | SNV: *PIK3CB*, *CCR6*, *CARD9*, *NLRP8*, *MAPK8* (1/1); *HDAC11*, *DEFB7* (0/1)<br>INS: *IRF6*, *TOLLIP*, *NUMBL* (1/1)<br>DEL: *SPINK5*, *JAM3*, *GSK3A* (1/1) |
| Tharparkar | SNV: *ADIPOQ*, *PIK3CB*, *CXCR1*, *CARD9*, *NLRP8*, *IGF1R*, *IRAK2* (1/1); *GSTP1* (0/1)<br>INS: *AP3B1*, *PIAS1*, *TLR3* (1/1)<br>DEL: *JAM3*, *GSK3A* (1/1)<br>Complex: *RPS6KA5*, *MTA1* (1/1) | SNV: *ADIPOQ*, *PIK3CB*, *CXCR1*, *CARD9*, *NLRP8*, *IGF1R*, *IRAK2* (1/1); *NLRP10*, *DDX21*, *GSTP1*, *DEFB7* (0/1)<br>INS: *AP3B1*, *RPS6KA5*, *MTA1*, *TLR3*, *TOLLIP*, *MMP12*, *NUMBL*, *TBKBP1* (1/1)<br>DEL: *JAM3*, *OUTB2*, *PPP4C*, *GSK3A* (1/1) |
| Sahiwal | SNV: *ADIPOQ*, *RFTN1*, *CXCR1*, *CARD9*, *NLRP8* (1/1); *NOD2*, *SIGLEC15* (0/1)<br>INS: *PIAS1*, *TRIM71*, *TOLLIP* (1/1)<br>DEL: *JAM3* (1/1) | SNV: *ADIPOQ*, *RFTN1*, *CXCR1*, *CARD9*, *NLRP8*, *SH2DIA* (1/1); *OAS2*, *NOD2*, *SIGLEC15*, *DEFB7*, *DMBT1* (0/1)<br>INS: *TNIP1*, *ITGB3*, *MTA1*, *TLR3*, *TOLLIP*, *NUMBL* (1/1)<br>DEL: *SPINK5*, *JAM3*, *GSK3A* (1/1) |
| Karan Fries | SNV: *ADIPOQ* (0/1)<br>INS: *AP3B1* (0/1), *NUMBL* (1/1)<br>DEL: *GSK3A*, *JAM3* (1/1) | SNV: *ZBP1* (1/1); *ADIPOQ*, *RFTN1*, *OAS2*, *CD300LF*, *MAPK8*, *DEFB7* (0/1)<br>INS: *AP3B1*, *IRF6*, *NFAT5*, *PIAS1*, *NUMBL*, *TBKBP1* (1/1)<br>DEL: *JAM3* (1/1) |
| Holstein Friesian | INS: *TOLLIP* (1/1)<br>DEL: *JAM3* (1/1) | INS: *TOLLIP*, *MMP12* (1/1)<br>DEL: *JAM3* (1/1) |
| All indicine (KG, GR, TP, SW) | SNV: *CARD9* (1/1), *NLRP8* (1/1)<br>DEL: *JAM3* (1/1) | SNV: *CARD9*, *NLRP8* (1/1)<br>INS: *NUMBL*, *TOLLIP* (1/1)<br>DEL: *JAM3*, *GSK3A* (1/1) |

In the indicine breeds, KG, GR, TP, and SW, we consistently observed high-impact homozygous SNVs (1/1) in two genes involved in PRR-mediated signaling and innate immune activation, *CARD9* and *NLRP8*. In KG, TP, and SW, a high impact homozygous SNV of *CXCR1* involved in chemokine signaling and immune cell recruitment, was seen. In TP, SW, and KF, homozygous SNVs of *ADIPOQ,* a gene involved in inflammatory regulation and in the activation of classical pathways of the complement system, was conserved. A variant in *CCR6*, involved in chemokine signaling and immune cell recruitment, was found only in KG and GR. Similarly, a SNV of *PIK3CB*, a gene mediating signaling in neutrophils at infection sites, was found in GR and TP. No high-impact SNVs were observed in the taurine breed (HF).

Some homozygous SNVs were breed-specific. In KG, we found *IGF1R,* associated with immune cell development, antibody production, and balancing inflammatory mediators, and *TRIM66*, involved in enhancing innate immune activation. GR had *MAPK8*, involved in inflammatory signaling, and TP had *IRAK2*, a key mediator of TLR and IL-1 receptor signaling. In SW, there were two breed specific homozygous variants: *RFTN1,* which participates in TLR as well as B-cell signaling, and *NOD2,* involved in activating innate immune response to bacterial and viral infection.

Heterozygous SNVs (0/1) in *C5*, a central component of complement activation, were observed in KG. Other breed-specific heterozygous SNVs included *DEFB7*, involved in antimicrobial defense, in GR, and *SIGLEC15*, involved in immune tolerance and prevention of excessive immune activation, in SW. TP had two breed-specific heterozygous SNVs: *DEFB7*, involved in antimicrobial defense, and *GSTP1,* involved in antioxidant defense and inflammatory modulation.

In KG, SW, and HF, homozygous insertions (1/1) were identified in *TOLLIP*, a key negative regulator of TLR-mediated inflammation. Homozygous insertions were also observed in *IRF6* (interferon-mediated antiviral signaling) in KG, *AP3B1* (intrinsic antiviral defense) and *TLR3* (PRR-mediated immune activation) in TP, and *NUMBL* (negative regulation of NF-kappaB signaling and inflammatory response) in KF. No high-impact insertions were observed in GR.

Homozygous deletions (1/1) in *JAM3*, essential for cell-cell adhesion and maintaining epithelial barrier integrity, were observed across all indicine, crossbred, and taurine breeds, suggesting conservation across the cattle groups. Deletions in *SPINK5* (regulation of NF-kappaB-mediated inflammation and antimicrobial defense) were shared between KG and GR. Deletions in *GSK3A* (involved in interferon-mediated antiviral response) were detected in GR and TP.

Overall, the indicine breeds, KG, GR, TP, and SW, showed a higher number of variants in immune-related genes than the crossbreed, KF and the taurine breed HF, potentially reflecting their adaptation to local environmental pressures and disease challenges.

Comparative functional clustering of genes with high-impact SNVs and SVs revealed key differences in immune functions across the cattle groups (Table 5). Indicine breeds exhibited greater functional variation across pathogen recognition, innate immune activation, inflammatory signaling, immune regulation, antiviral signaling, complement activation and chemokine signaling. In TP, functional differences were greater, with variants affecting pathogen sensing (*TLR3*), key innate immune mediators (*CARD9, NLRP8, IRAK2*), signaling hubs (*PIK3CB, JAK1, BCL10*), antiviral signaling (*GSK3A, AP3B1*), and complement components (*C8A, C8B*), suggesting strong pathogen-driven selection. In contrast, the crossbreed KF had moderate functional changes, mainly in inflammatory regulation and dampening interferon signal, reflecting partial retention of indicine immune traits. HF showed minimal immune functional variation, with variants mainly confined to regulating inflammation and maintaining the epithelial barrier. Collectively, these findings highlight key genomic features underlying resistance to disease and environmental pressures in indicine cattle, particularly enriched in TP.

**Table 5.** Functional clustering of innate immune genes carrying high-impact small variants and structural variants across the six cattle breeds.

| Functions of genes and gene clusters | Kangayam | Tharparkar | Gir | Sahiwal | Karan | Holstein |
| --- | --- | --- | --- | --- | --- | --- |
| Barrier integrity maintenance and epithelial immune defence | *JAM3*,<br>*PCDH1* | *JAM3* | *JAM3* | *JAM3* | *JAM3* | *JAM3* |
| Antimicrobial defence | - | - | *DEFB7* | - | - | - |
| Pathogen recognition and innate immune activation | *CARD9*,<br>*NLRP8* | *CARD9*,<br>*NLRP8*,<br>*TLR3*, *IRAK2* | *CARD9*,<br>*NLRP8*,<br>*PTX3* | *CARD9*,<br>*NLRP8*,<br>*NOD2*,<br>*RFTN1* | - | - |
| Complement activation and regulation | *C5* | *C8A*, *C8B*,<br>*ADIPOQ* | *PTX3* | *ADIPOQ* | *ADIPOQ* | - |
| Antiviral signaling and interferon modulation | *IRF6*, *G3BP1* | *GSK3A*,<br>*AP3B1* | *GSK3A*,<br>*DHX36*,<br>*PLSCR1*,<br>*TRIM42* | - | - | - |
| Negative regulation of antiviral and interferon signaling | - | *DR1* | - | - | *CRLF3* | - |
| Inflammatory signaling and immune cell activation | - | *PIK3CB*, *JUN*,<br>*JAK1*, *BCL10*,<br>*CD53*, *GNAI3* | *MAPK8*,<br>*PIK3CB* | - | *TP73* | - |
| Chemokine signaling and immune cell recruitment | *CXCR1*,<br>*CCR6* | *CXCR1* | *CCR6*,<br>*P2RY14* | *CXCR1* | - | - |
| Positive regulation of innate immune activation | *TRIM66* | - | - | - | - | - |
| Regulation of inflammatory response | *CAMK2A*,<br>*EDIL3*,<br>*RGMB*,<br>*HAVCR2*,<br>*CNOT8* | *ADIPOQ* | *SIAH2* | *ADIPOQ* | *ADIPOQ*,<br>*NUMBL* | - |
| Redox regulation in immune signaling | *GLRX* | *GSTP1* | - | - | - | - |
| Negative regulation of inflammatory response | *TOLLIP*,<br>*STAMBP*,<br>*SPINK5* | *STAMBP* | *SPINK5* | *TOLLIP*,<br>*STAMBP* | - | *TOLLIP* |
| Immune homeostasis and resolution of inflammation | *NR3C1*,<br>*TNIP1*,<br>*TNIP3*,<br>*PPARGC1B* | *TNIP3* | - | - | - | - |
| Apoptosis mediated immune regulations | - | - | *TNFSF10* | *MEGF10* | - | - |
| Immune cell survival and immune balance | *IGF1R* | *IGF1R* | - | - | - | - |
| Antigen processing and modulation of innate immunity | *ERAP1* | - | - | - | - | - |
| Linking innate and humoral immunity | *IL12B* | - | *IL12A* | - | - | - |
| Immune tolerance and regulatory signaling | *ADRB2* | - | *SIAH2* | *SIGLEC15* | - | - |

### Indicine-specific high-impact variants in *CARD9* and *NLRP8*

Across all four indicine breeds (KG, GR, TP, and SW), FreeBayes identified commonly shared variants including SNVs in 1,647 genes, deletions in 36 genes, complex variants in 32 genes, MNVs in 25 genes, and insertions in 23 genes. In contrast, GATK detected 32,096 non-immune and 973 immune-related genes (629 innate) with shared variants, comprising 955 immune genes with SNVs, 203 with insertions, and 176 with deletions.

Among these, FreeBayes detected high-impact variants in five immune genes (including three innate immune genes), while GATK identified 10 immune genes (including six innate immune genes) (Table 4). Both callers consistently captured high-impact variants in three innate immune genes, *CARD9* (rs132699263), where a T→C substitution on chromosome 11 resulted in stop-loss mutation, *NLRP8* (rs42734179) with a T→C splice-donor disruption on chromosome 18, and *JAM3* (no rsID match) with a GA→G frameshift deletion on chromosome 29 (Table 4).

Comparative analysis showed that the high-impact variants, *CARD9* and *NLRP8*, were present in all indicine breeds but absent in KF and HF, suggesting indicine-specific adaptive signatures. Variant calling with pooled-sample settings using FreeBayes and GATK (ploidy=12) revealed that variants in *CARD9* and *NLRP8* were fixed across all alleles (1/1/1/1/1/1/1/1/1/1/1/1) in each pool. This indicates that all six animals in each pool carried these variants relative to the reference genome.

*CARD9* encodes for a caspase recruitment domain containing the adaptor protein (CARD9) that plays a key role in innate and adaptive immunity. The gene mediates signaling downstream of PRRs, including C-type lectin receptors (CLRs), nucleotide-binding oligomerization domain (NOD) receptors, and TLRs. Through these signaling, CARD9 induces NF-κB and MAPK activation, triggering inflammatory cytokine responses that are essential for effective defense against microbial infections [47]. *NLRP8* (NOD-like receptor family pyrin domain containing 8) belongs to the family of intracellular PRRs. The NLR protein family is a recently recognized class of innate immune molecules in animals [48]. The two genes, *CARD9* and *NLRP8*, could be potential immune markers associated with disease resistance in indicine breeds. However, experimental investigations are required to confirm their functional significance in immune response and disease resilience.

### Reduced nucleotide diversity reflects distinct population structure in indicine breeds

Nucleotide diversity (π) measures genetic variations within populations and reflects historical breeding patterns, population size, and selection pressure [49]. Higher π values indicate greater genetic heterogeneity, whereas lower π values suggest reduced diversity due to factors such as selection, inbreeding, or population bottlenecks. Our analysis with SNV data revealed that HF and KF exhibited higher nucleotide diversity than the indicine breeds, KG, GR, TP, and SW (Figure 6A). This increased diversity in HF reflects a broader genetic background, admixture, artificial selection for dairy traits and the mixing of animals from many different genetic backgrounds through breeding programs [49]. KF displayed intermediate yet elevated diversity due to its indicine-taurine ancestry [38]. In contrast, indicine breeds showed lower genetic diversity, which likely reflects long-term adaptation to specific local environments, smaller effective population sizes, and limited selective breeding, resulting in more homogeneous genomic backgrounds [50].

**Figure 6.**
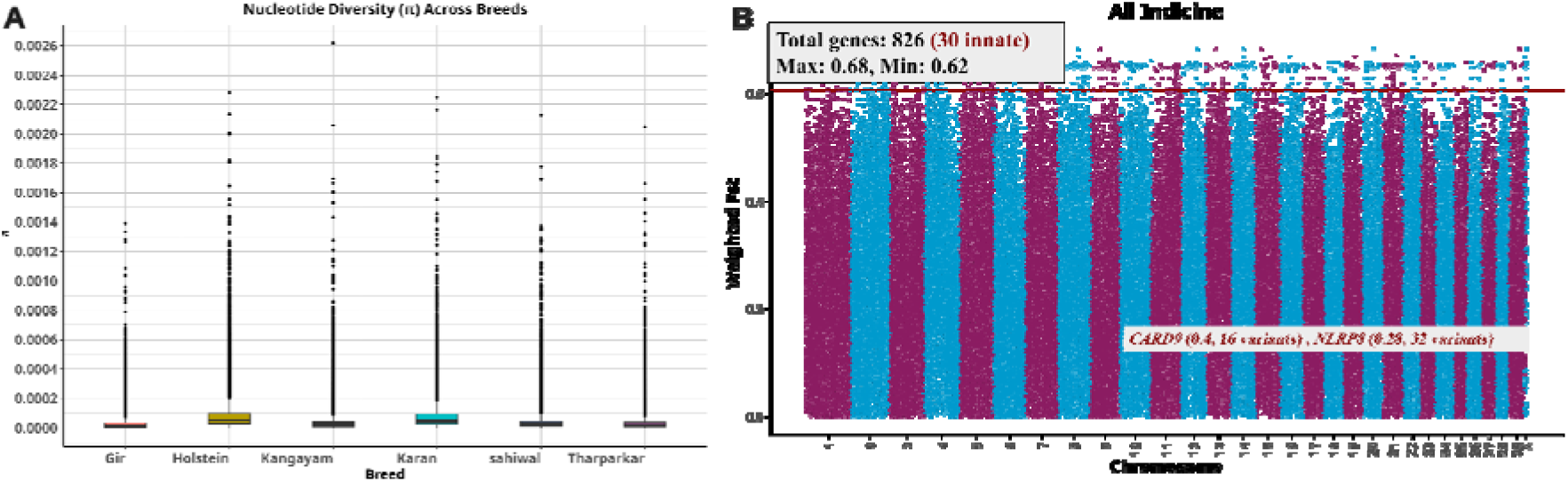
Nucleotide diversity (π) and F_ST_ analysis across breeds based on SNV genotype. (A) Nucleotide diversity (π) analysis of the six cattle breeds revealed that KF and HF had higher nucleotide diversity than indicine breeds. (B) Pairwise F_ST_ values, calculated based on the Weir and Cockerham method, between indicine (KG, GR, TP, and SW), crossbred (KF), and exotic (HF) breeds. The red line represents the top 1% threshold.

### Genetic differentiation of immune-related genes revealed by F_ST_ analysis

The F_ST_ analysis, across indicine (KG, TP, SW, and GR), exotic (HF) and crossbred (KF) breeds, revealed genetic differentiation among the three major cattle groups (Figure 6B). Breed-specific F_ST_ comparison, where each indicine breed was individually compared against the exotic as well as the crossbreed groups, highlighted breed-specific patterns of genomic divergence (Figure 7A–7D).

**Figure 7.**
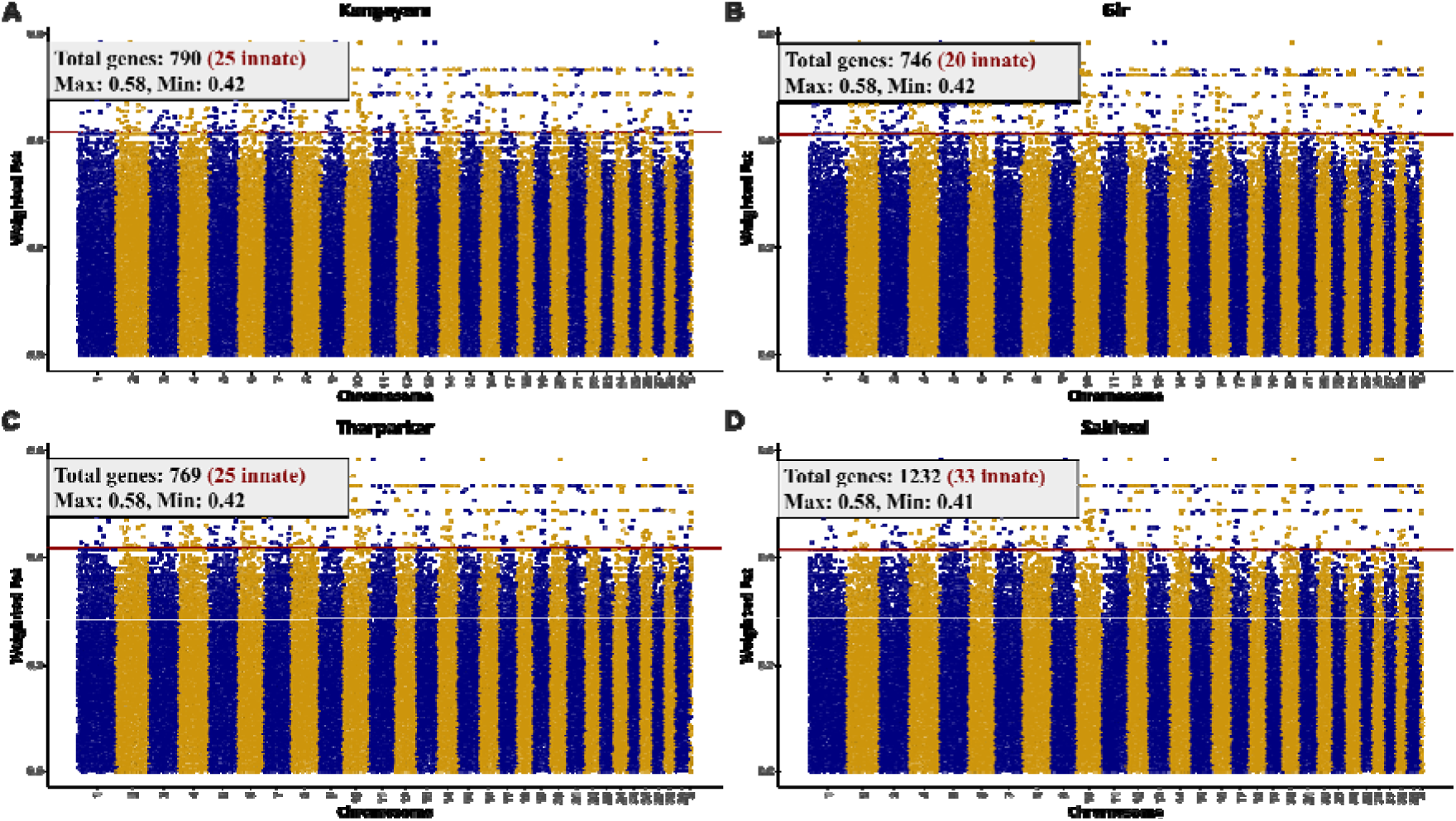
F_ST_ analysis of the four indicine breeds, the crossbreed and the exotic breeds. **Breed-**specific pairwise F_ST_ analysis of KG (A), GR (B), TP (C), and SW (D) with KF and HF. The red line represents the top 1% threshold.

Several genomic regions showed strong differentiation (groupwise F_ST_ > 0.6; breed-specific F_ST_ > 0.4) in the top 1% category. Group-wise comparison revealed 826 highly differentiated genes, of which 30 were innate immune genes, highlighting strong immune-driven divergence among cattle groups. Several innate immune genes exhibited high F_ST_ values despite having only a small number of SNVs within the corresponding genomic windows. Among the 30 innate immune genes, 15 genes showed high F_ST_ values with a single SNV per window, including, *E2F1* (0.66), *CCDC88A* (0.65), *TRADD* (0.64), *ATF2* (0.63), and *GAB1* (0.63).

These genes are known to play critical roles in innate immune functions. For example, *TRADD* is involved in antiviral response and programmed cell death signaling, while *GAB1* activates TLR3/4-and RIG-I-triggers innate responses and induces the type-I interferon. In addition, SCN5A functions as a pathogen sensor that regulates ATF2-mediated transcription to initiate antiviral signaling [51]. *E2F1* is an important transcription factor that is primarily involved in regulating innate immune and inflammatory response, and apoptosis [52]. *CCDC88A* encodes girdin, also known as Gα-interacting vesicle-associated protein (GIV), which interacts with the cytosolic pattern recognition receptor NOD2 in macrophages. This interaction is essential for balancing antimicrobial defense and anti-inflammatory responses by suppressing NF-κB–dependent inflammatory signaling and promoting phagolysosomal fusion and bacterial clearance, preventing excessive inflammation and maintaining effective microbial control [53].

Nine innate genes showed high F_ST_ values with two SNVs, including *EDIL3* (0.66), *ADAM10* (0.65), *CD209* (0.65), and *TP63* (0.63). *TP63* regulates interferon-gamma-mediated signaling and apoptosis pathways. *CD209* mediates pathogen recognition with initiation of innate as well as adaptive immune responses [54]. *ADAM10,* a key regulator of lymphocyte development, including T-cell and marginal zone B-cell differentiation, is essential for the proper positioning of T and B cells during immune responses [55]. Additionally, four genes (*SREBF2, TNK1, MAP3K12,* and *RF2BP1*) showed high F_ST_ score with three SNVs and one gene (*TP53*) with four SNVs within the window.

The innate gene, *C1R*, showed a high F_ST_ value (0.63) with seven SNVs within the windows. *C1R* encodes complement component C1r, a serine protease of the C1 complex, the first component of the classical complement pathway that initiates the proteolytic cascade upon activation [56]. Indicine-specific innate genes, such as *CARD9* (F_ST_ = 0.4; 16 variants) and *NLRP8* (F_ST_ = 0.28; 32 variants), absent in KF and HF, were identified by FreeBayes and GATK, highlighting potential indicine-specific immune adaptations and strong selective pressure acting on this locus.

Breed-specific F_ST_ analysis (top 1%; F_ST_ > 0.4) also revealed distinct patterns of innate immune divergence of the indicine breeds from the crossbreed and the taurine breed. In KG (Figure 7A), 790 genes, including 25 innate immune genes, were identified within the top 1% F_ST_ region. GR (Figure 7B) showed 746 genes, including 20 innate genes, while TP (Figure 7C) exhibited 769 genes, including 25 innate genes. The highest level of divergence was observed across multiple genes in SW (Figure 7D), with 1232 genes, including 33 innate genes, observed in the top 1% F_ST_ regions.

### Integrating relative (F_ST_) and absolute divergence (d_XY_) identifies immune signatures in indicine breeds

To identify genomic regions representing true genetic divergence, the top 1% F_ST_ windows were correlated with d_XY_ results for each breed. This integrative analysis identified a subset of innate immune genes showing consistently high divergence across both metrics (F_ST_ and d_XY_ ≈ 0.5), highlighting potential breed-specific immune signatures.

In KG, seven innate genes, *E2F1*, *SREBF2, NLRP3, EDIL3, CYLD, ATF2,* and *IL1A*, showed strong divergence. *SREBF2* encodes the transcription factor, SREBP2, which regulates cholesterol homeostasis and contributes to leukocyte innate and adaptive immune responses by upregulating cholesterol flux and activating pro-inflammatory genes such as *NLRP3, NOX2,* and *IL8* [57]. Inhibiting SREBP2 has been shown to reduce lipid metabolism essential for viral replication, limiting viral load [58, 59]. *NLRP3* functions as an inflammasome sensor of pathogen- and damage-associated molecular patterns (PAMPs and DAMPs), triggering pro-inflammatory cytokine (IL-1β and IL-18) release and linking innate to adaptive immune responses [60]. *CYLD*, a deubiquitinase that removes K63-linked ubiquitin chains, is essential for antiviral defense. Its loss impairs type I IFN production, increases susceptibility to viral infection, making it a key regulator of antiviral innate immunity [61]. *IL1A* is known to mediate protective immune response during bacterial and viral infection [62].

In GR, six innate genes, *E2F1, EDIL3, SREBF2, BECN1, CYLD,* and *NLRP3*, exhibited high divergence in both analyses. *BECN1* (Beclin 1) is a highly conserved autophagy regulator that plays an important role in innate immunity. It modulates apoptosis and phagocytosis, suppresses NF-κB–dependent inflammation, and interacts with immune signaling pathways. A previous study reported that silencing *BECN1* enhances antiviral immunity and increases survival rate [63].

TP showed the highest number of strongly divergent innate genes, with ten genes identified: *NFKB2, BECN1, SREBF2, CYLD, E2F1, NLRP3, EDIL3, ATF2, CD53,* and *IL1A. NFKB2* encodes the p100/p52 subunit of the NF-κB family and mediates non-canonical NF-κB signaling, essential for regulating lymphocyte and macrophage functions during immune and inflammatory responses [64].

In SW, seven innate genes, *EDIL3, RORA, SREBF2, BECN1, CYLD, E2F1,* and *ATF2*, showed strong divergence, supported by both metrics. *RORA* encodes retinoid-related orphan receptor α (RORα), a nuclear receptor and transcription factor involved in circadian rhythm, cholesterol metabolism, and immune regulation. RORα controls inflammation through the NF-κB pathway. Genetic variation in *RORA* influences the strength of trained immunity responses [65].

Across all indicine breeds, *SREBF2, E2F1, EDIL3,* and *CYLD* consistently showed high divergence in both F_ST_ and d_XY_ analyses, indicating shared immune-related selective pressures. While F_ST_ captures relative genetic differentiation across multiple populations through weighted estimates, d_XY_ provides absolute divergence between two populations. Integrating both metrics revealed that several innate immune genes exhibited high divergence between indicine and taurine breeds but comparatively lower divergence between indicine breeds and the crossbreed.

*TRADD* showed high differentiation in F_ST_ analysis, while d_XY_ revealed greater divergence between indicine and crossbreed populations and lower divergence between indicine and taurine breeds. This pattern was consistently observed across all indicine breeds. Similarly, *MALT1* displayed high differentiation in the SW-specific F_ST_ analysis, while d_XY_ indicated higher divergence between SW and KF and lower divergence between SW and HF. *MALT1* is a key regulator of both innate and adaptive immune signaling and plays an essential role in host defense against microbial infections [66, 67].

The higher absolute divergence observed between indicine and crossbreed populations, together with the lower divergence between indicine and taurine breeds, indicates that these immune genes have likely been modified by recent introgression and selective breeding in crossbreed cattle rather than reflecting deep ancestral divergence. This pattern underscores the influence of admixture and artificial selection on the evolution of immune-related genes.

### Selective sweep detection by RAiSD

Genome-wide selection scans, using the μ statistic in RAiSD, identified multiple positively selected regions across 29 autosomes and X chromosomes in all six cattle breeds (Figure 8). The μ statistic is the selective sweep score, where a high μ value indicates strong evidence of a recent selective sweep. It represents overall sweep signature strength, with peaks indicating candidate regions under positive selection. RAiSD integrates signals, such as reduced diversity, shifts in allele frequency, and increased linkage disequilibrium, making it effective for detecting recent adaptive events. Figure 8 indicates that selection signals were unevenly distributed across chromosomes in all six breeds. Chromosome 1 showed the highest number of signals, likely reflecting its larger size (158 Mb) and greater gene content compared with other chromosomes. In addition, several chromosomes showed breed-specific enrichment of selective sweep signals.

**Figure 8.**
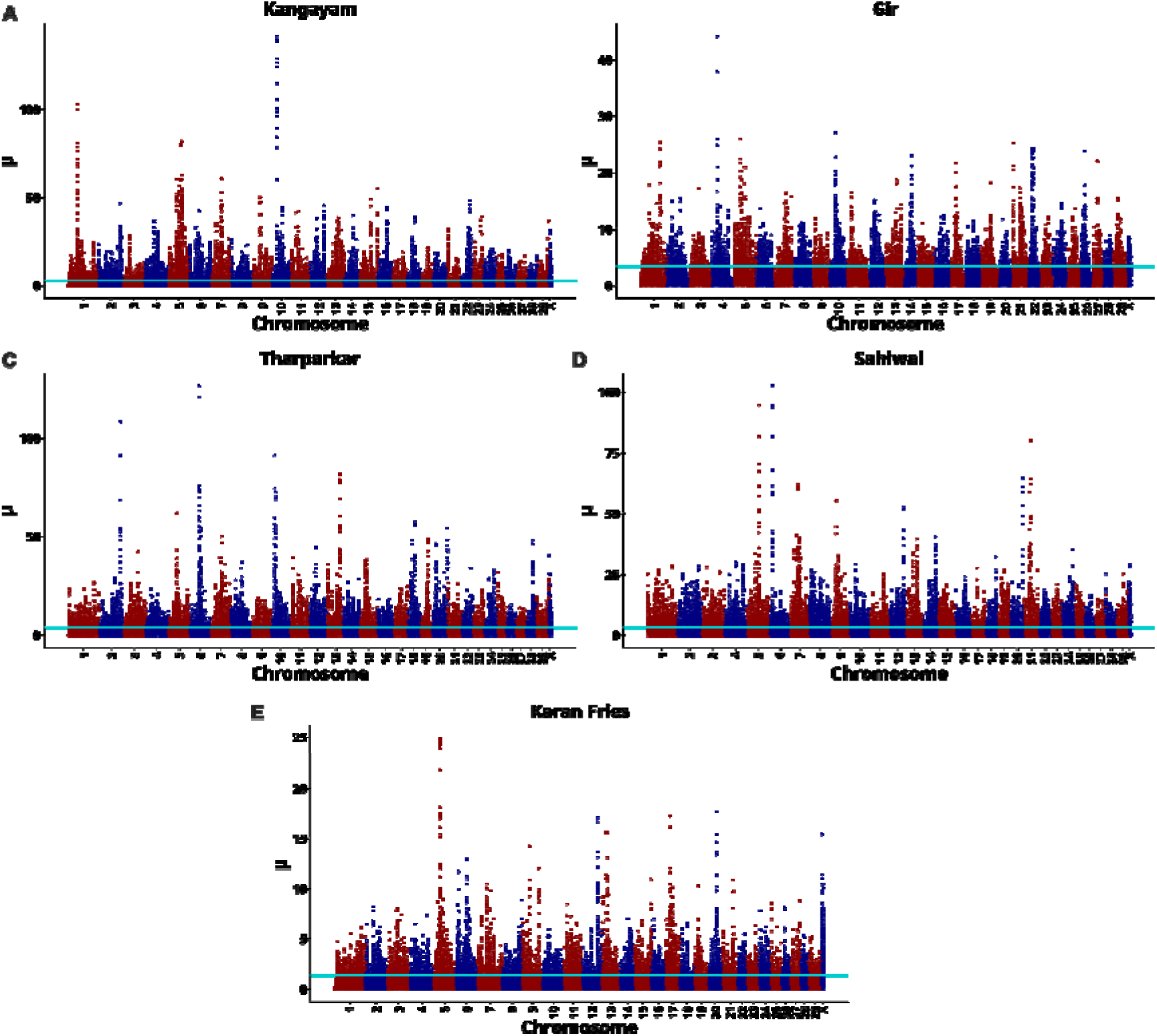
Manhattan plot showing RAiSD selective sweep signals across the genome of the six cattle breeds. The x-axis represents genomic position ordered by chromosome (chr1-chr29 and X), while the y-axis shows the RAiSD µ statistic, with higher values indicating stronger evidence of positive selection. The cyan horizontal line indicates the top 1% threshold of selective sweep signals. The panel shows µ statistics for (A) KG, (B) GR, (C) TP, (D) SW, and (E) KF.

In KG (Figure 8A), a total of 2543 genes fell within the top 1% μ score, including 142 immune-related genes (107 innate), with the innate gene, *IRAK3* (important modulator of inflammation in innate immunity [68]), producing a strong signal (μ = 45.46). GR (Figure 8B) showed 1366 candidate genes, including 72 immune-related genes (55 innate), with *RNF41* (promotes anti-inflammatory defense [69] and TRIF-dependent production of type I interferon [70]) as the top innate gene (μ = 14.59). TP (Figure 8C) displayed 2486 candidate genes, including 121 immune-related genes (89 innate), with the innate gene, *ITCH1* (regulates the interplay between innate and adaptive immune response [71] and dampens inflammatory signaling [72]), showing the highest μ value (69.39). SW (Figure 8D) displayed 2633 genes, including 131 that were immune-related (92 innate), with *RCAN1* (regulates immune and inflammatory responses during infection and limits excessive cytokine production [73]) (μ = 25.35) showing the highest signal across the innate gene.

KF (Figure 8E) had 1821 genes, including 91 immune-related (70 innate) genes, with *ANO6* (essential for immune defence, facilitates bacterial phagocytosis [74]) (μ = 24.87) as the top innate candidate. When the data from the indicine breeds, KG, GR, TP, and SW, were combined (Supplementary Figure 3), 465 genes fell within the top 1%, including 24 immune-related (15 innate) genes. However, the μ values were lower (maximum 1.40 × 10^−10^, minimum 5.71 × 10^−11^), suggesting that selective sweeps are mostly breed-specific, likely driven by local environmental and pathogenic pressures. Overall, the enrichment of immune and innate immune genes under selection supports the idea that pathogen-driven selection has played a major role in shaping genetic adaptation in cattle, consistent with earlier findings in livestock species [75–77].

### Enrichment analysis of genes within selective sweep regions

Across the indicine breeds, KG, GR, TP, and SW, the functional enrichment analysis of genes located within the top 1% of selective sweep regions revealed significant KEGG pathways and biological processes related to gene ontology (GO) (p-value < 0.05) (Supplementary Table 5 and Supplementary Table 6). These enriched terms were broadly related to adaptation, reproduction, growth, immune function, and production traits, highlighting the biological relevance of selective sweeps in indicine breeds.

#### Kangayam (KG)

A total of 67 KEGG pathways were significantly enriched (Supplementary Table 5). Among the top 1% sweep regions, many genes were related to MAPK (57 genes), Rap1 (45 genes), and cAMP signaling (45 genes). Pathways related to endocytosis (43 genes), T cell receptor signaling (24 genes), inflammatory mediator regulation of TRP channels (22 genes), C-type lectin receptor signaling (19 genes), B cell receptor signaling (18 genes), Fc gamma R-mediated phagocytosis (18 genes), and those involved in the bacterial invasion of epithelial cells (15 genes) were also enriched in KG.

In GO biological processes (157 terms), the most enriched were the regulation of transcription by RNA polymerase II (163 genes), signal transduction (85 genes), protein phosphorylation (63 genes) and protein ubiquitination (49 genes). Immune processes included endocytosis (26 genes), T-cell receptor signaling (15 genes), negative regulation of canonical NF-kappaB signal transduction (12 genes), T cell proliferation (9 genes), and positive regulation of T-helper 1 cytokine production (4 genes) (Supplementary Table 6).

#### Gir (GR)

GR showed enrichment in 47 KEGG pathways (Supplementary Table 5). Pathways with the highest gene counts included pathways in cancer (46 genes), human papillomavirus infection (30 genes), MAPK (28 genes), and Rap 1 signaling (26 genes). Immune-related enrichment was observed in human papillomavirus infection and the C-type lectin receptor signaling pathway (13 genes). As with KG, GO biological process analysis (126 terms) showed major enrichment in the regulation of transcription by RNA polymerase II (106 genes), signal transduction (54 genes), and protein phosphorylation (34 genes) (Supplementary Table 6).

#### Tharparkar (TP)

TP showed enrichment in 62 KEGG pathways, with the highest gene counts observed in metabolic pathways (185 genes), pathways in cancer (71 genes), MAPK signaling (44 genes) and the oxytocin signaling pathway (32 genes) (Supplementary Table 5). Immunity-related pathways enriched in TP included T cell receptor signaling (23 genes), inflammatory mediator regulation of TRP channels (18 genes), and Fc epsilon RI signaling (13 genes). GO analysis identified 169 enriched biological processes, with major categories similar to those in KG, such as the regulation of transcription by RNA polymerase II (186 genes), signal transduction (76 genes), protein phosphorylation (60 genes), protein ubiquitination (46 genes), and the apoptotic process (44 genes). Immune-related GO terms included endocytosis (22 genes), T cell receptor signaling (17 genes), negative regulation of toll-like receptor signaling (5 genes), and regulation of TNF production (4 genes) (Supplementary Table 6).

#### Sahiwal (SW)

In Sahiwal, 74 KEGG pathways were significantly enriched. The most prominently enriched were pathways in cancer (74 genes), cAMP signaling (55 genes), PI3K-Akt signaling (54 genes), MAPK signaling (45 genes) and the immunity-related pathway involved in the bacterial invasion of epithelial cells (16 genes) (Supplementary Table 5). Significantly enriched GO biological process terms came to 144. As with KG, TP, and GR, enrichment was primarily observed in the regulation of transcription by RNA polymerase II (164 genes), signal transduction (86 genes), protein phosphorylation (66 genes), and endocytosis (27 genes) (Supplementary Table 6).

### Identification of potential innate immune genes in indicine breeds and their association with gene expression

Potential innate immune genes were identified by integrating genes with SVs, and CNVs from DELLY, small variants (SNVs and InDels) from FreeBayes and GATK, as well as candidate genes within the top 1% of F_ST_ and RAiSD selective sweep regions. Of a total of 1226 curated innate immune genes in InnateDB, this integrative filtering yielded non-redundant sets of 319 genes in KG, 215 in GR, 304 in TP, 292 in SW, 144 in KF, and 71 in HF. Compared to KF and HF, indicine breeds consistently showed a higher number of potential innate immune genes, reflecting stronger historical selection and adaptation of innate immunity.

To evaluate the functional relevance of genes with variants, the identified innate immune genes were correlated with previously published PBMC differential expression profiles from healthy indicine breeds (GR, TP, SW) and a crossbreed (KF) [32]. This correlation revealed several innate immune genes carrying genomic variants, located within F_ST_ and selective sweep regions, with distinct basal expression patterns under non-infected conditions (Table 6), indicating inherent differences in baseline immune responsiveness among breeds.

**Table 6:** Correlation of innate immune genes carrying genomic variants with differential expression between indicine breeds (GR, TP, and SW) and a crossbreed (KF). Cells represented in bold, with a single asterisk (*), indicate genes identified by F_ST_ and differential gene expression analyses. Cells in bold,with a double asterisk (**), indicate genes identified by selective sweep and differential gene expression analyses. Genes common to both are in bold with the $ symbol. Cells in bold, with a triple asterisk (***), indicate genes having missense variants and are differentially expressed.

| Upregulated genes |  |  | Downregulated genes |  |  |
| --- | --- | --- | --- | --- | --- |
| Gir | Tharparkar | Sahiwal | Gir | Tharparkar | Sahiwal |
| **PCDH1 \*** | *PCDH1* | **PCDH1 \$** | *IFITM1* | *IFITM1* | *IFITM1* |
| **LRP1B \$** | **LRP1B \$** | **LRP1B \$** | **IGDCC3 \*** | **IGDCC3 \*\*\*** | *IL32* |
| *CD300E* | *CD300E* | *PDGFA* | *ADAM11* | *ADAM11* | *ADAM11* |
| *CCL8* | *CCL8* | *CCL8* |  | **TYRO3 \*\*** | **TYRO3 \*\*** |
| **WNT4 \*** | *WNT4* | *WNT4* |  | *F2RL3* | *FKBP14* |
| **OAS2 \*\*\*** | *FOS* | *SOX4* |  | **SYT7 \*\*** | *STEAP2* |
|  | *PDGFA* | *CEBPE* |  | *TREML1* | *CDH2* |
|  | **CELSR1 \*\*\*** | **CELSR1 \*\*\*** |  | **SPOCK1 \*\*** | **SPOCK1 \*\*** |
|  | **RNF128 \*\*** | *DNASE1L3* |  | **LTC4S \*\*\*** | *ADRA2A* |
|  | *PRKRA* | *HPGD* |  | *CLU* | *SYCP2* |
|  | *PLAUR* | *ACHE* |  | *STEAP2* | *DDR1* |
|  | *CD34* | *PLA2G7* |  | *TRIM7* | *TRIM7* |
|  | *ACHE* | *RRAD* |  |  | *C3AR1* |
|  | *MSX1* | *LMO7* |  |  | *GSDMA* |
|  | *COL11A1* | **LAMA4 \*** |  |  | **CR2 \*** |
|  | **SEMA6A \*** | *ITGB6* |  |  |  |
|  | *FN1* | **TNNT3 \*\*** |  |  |  |
|  | *SMAD7* | **RETN \*\*\*** |  |  |  |
|  | *KCNH8* | *TACR1* |  |  |  |
|  | *DGKG* | *MMP3* |  |  |  |
|  | *NR4A3* | **NR4A3 \*\*\*** |  |  |  |
|  | **NLRP13 \*\*\*** | *S100A7* |  |  |  |
|  |  | *DEFB1* |  |  |  |
|  |  | *ISG15* |  |  |  |

Compared with KF, across all indicine breeds, we observed consistent upregulation of *LRP1B*, identified in both F_ST_ and selective sweep analysis. *LRP1B* encodes a low-density lipoprotein receptor involved in immunity regulation through the suppression of hedgehog signalling [78]. *LRP1B* has been previously reported within selection signatures of Vrindavani cattle, where it was targeted for traits related to productivity and environmental adaptation [79]. This gene has also been associated with milk yield and somatic cell count [80]. The consistent upregulation of this gene across indicine breeds indicates that it provides better immunity in indicine breeds than in KF [36].

Similarly, protocadherin-1 *(PCDH1)* exhibited strong upregulation (>9-fold) in indicine breeds [32], suggesting better immunity compared to KF [36]. *PCDH1*, identified in F_ST_ and in the top 1% of selective sweep regions, had missense variants in SW and within F_ST_ regions in GR. No *PCDH1* variants were identified in TP. *PCDH1* is a glucocorticoid regulator that plays a key role in maintaining cell–cell adhesion and epithelial barrier integrity, a critical first line of innate defence against pathogens [81]. Disruption of epithelial tight junctions facilitates pathogen invasion and promotes viral, bacterial, and fungal infections [82], highlighting the functional importance of *PCDH1* variation and expression differences.

In contrast, *SPOCK1* and *TYRO3,* both located within the top 1% of selective sweep regions, were consistently downregulated in TP and SW. In addition, several innate immune genes having missense variants, identified by FreeBayes and GATK, showed concordant changes in gene expression. These included *OAS2* in GR, *NLRP13, CELSR1, CD34, LTC4S, IGDCC3,* and *TYRO3* in TP, and *CELSR1, RETN, PCDH1, NR4A3, SPOCK1, TYRO3, CR2* in SW. The overlap of genomic variants, selective sweep signals, and differential basal expression supports the functional relevance of these genes in shaping breed-specific innate immunity. The functional roles of these genes could provide deeper insights into disease resistance in indicine breeds. Future studies need to integrate transcriptome data from infected or challenged conditions to investigate how the genes carrying genomic variants respond during active immune stimulation.

## Conclusion

We provide a detailed, genome-wide view of immune-related genetic variation across indicine, taurine, and crossbreed cattle, offering insights into the genomic basis of disease resilience in breeds adapted to the tropics. By combining whole-genome sequencing with population structure analysis, structural and small variant detection, selective sweep detection, and basal gene expression profiling, compared to the HF and KF breeds, in indicine cattle, we identified a distinctive repertoire of immune-associated variants. Multiple innate immune genes involved in pathogen sensing, inflammatory signaling, antigen processing, and host defense were repeatedly identified across independent analyses. High-impact variants in the innate immune genes, *CARD9* and *NLRP8*, unique to indicine breeds, were absent in taurine and crossbred cattle. The integration of genomic variation with transcriptional responses strengthens the biological relevance of these candidate markers, linking genetic signatures to functional immune responsiveness.

The genomic resources and candidate gene sets generated in this study provide a foundation for future investigations into the molecular mechanisms underlying disease resistance in cattle. The data can be used to prioritise functional validation studies, explore genotype–phenotype relationships across diverse diseases, and assess how immune-associated variants interact with environmental stressors such as heat and pathogens. The results underscore the value of preserving and strategically incorporating indicine immune diversity into crossbreeding and genomic selection programs. Leveraging such immune-adaptive genomic information is critical for the sustainable development of productive cattle populations with resilience against diseases and changes in climate.

## List of Abbreviations

CPG -: Copy number gain
CPL -: Copy number loss
CNV -: Copy number variant GO - Gene ontology
GR -: Gir
HF -: Holstein Friesian
InDel -: Insertion and deletion
KG -: Kangayam
KF -: Karan Fries
L1 -: Location 1
L2 -: Location 2
L3 -: Location 3
MAPQ -: Mapping quality
MAF -: Minor allele frequency
MNV -: Multi-nucleotide variant
PBMC -: Peripheral blood mononuclear cell
PCA -: Principal component analysis
PRR -: Pattern recognition receptors
SW -: Sahiwal
SNV -: Single-nucleotide variant
SV -: Structural variant
TP -: Tharparkar
TLR -: Toll-like receptor

## Acknowledgement

The authors thank the ICAR-National Agricultural Science Fund for financial support, SASTRA Deemed to be University, and ICAR-National Dairy Research Institute for infrastructural support and Dr. Ashutosh Vats for the gene expression experiments. The authors thank Madhu and Gita for critical discussions and revisions of the manuscript.

## Data Availability

Raw variant data generated using DELLY and FreeBayes have been deposited in the European Variant Archive (EVA) under the project accession PRJEB88461 (https://www.ebi.ac.uk/eva/?eva-study=PRJEB88461). All additional supporting data are provided in the Supplementary Materials accompanying this article.

