## Supplementary Figures for "Integrative Whole-Genome Analysis reveals Genomic Signatures of Innate Immunity in Indicine Cattle"

**Supplementary Figure 1: The detailed representation of the sample preparation.** The samples including blood and semen were collected from six healthy cattle breeds from three different locations for each breed. At each location, samples from six different animals of the same breed were pooled, resulting 18 pooled samples (1 pooled sample of 6 animals x 6 breeds x 3 locations) in total. These pooled sample were utilized for whole genome sequencing.

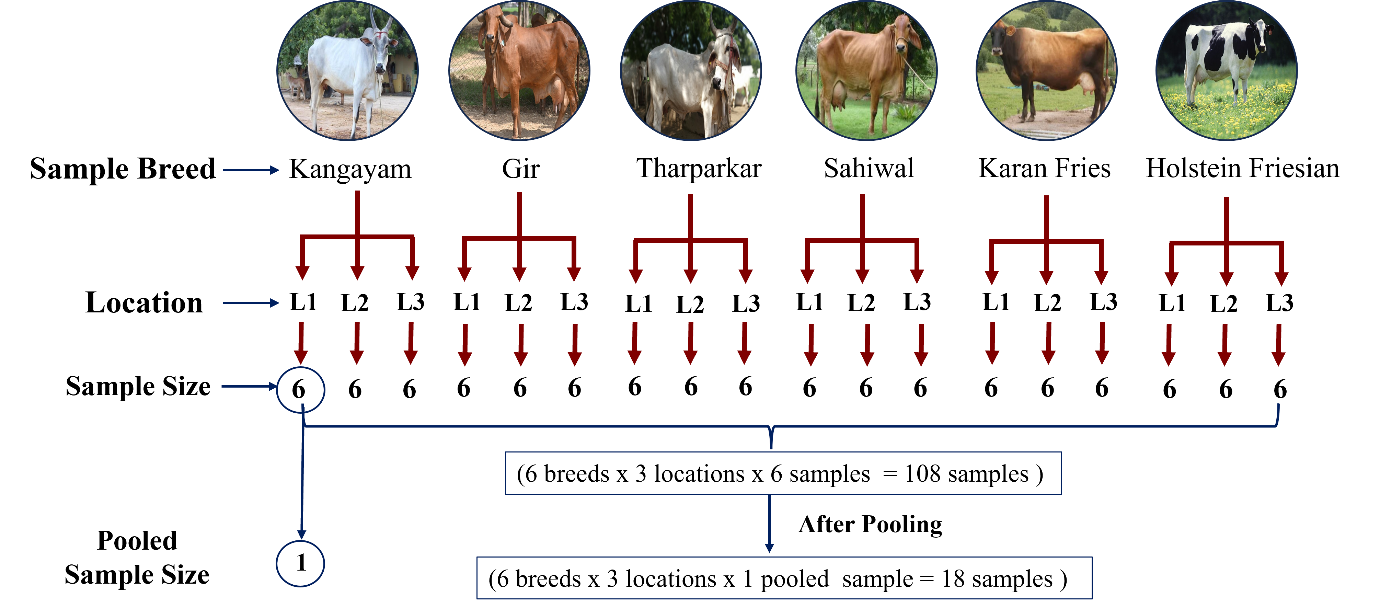

**Supplementary Figure 2: The alignment statistics and mapping quality score for 18 pooled samples**

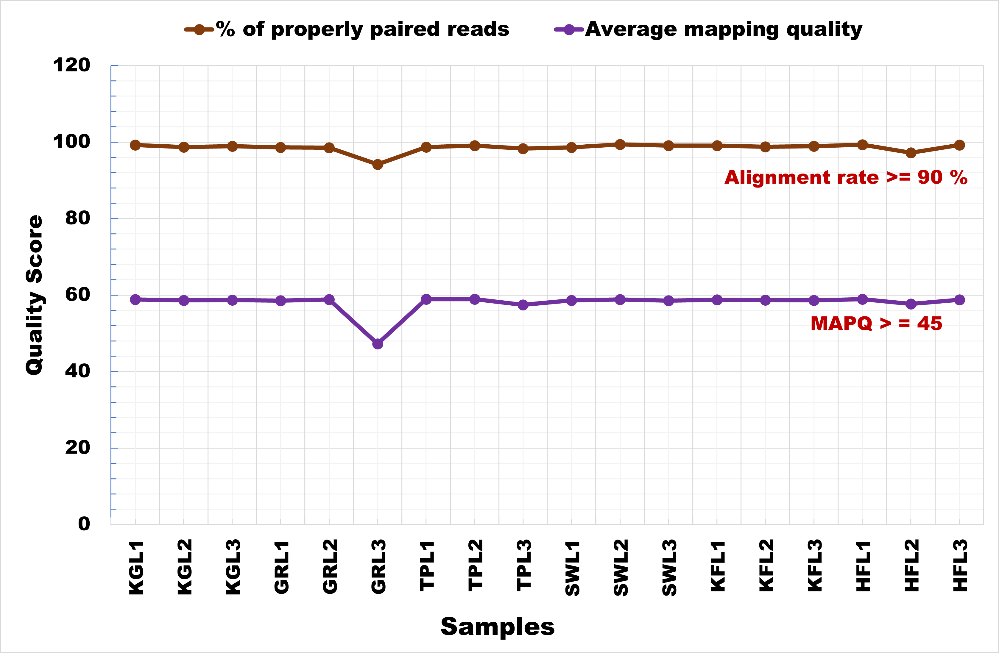

**Supplementary Figure 3: µ statistics for RAiSD analysis in indicine breeds.**

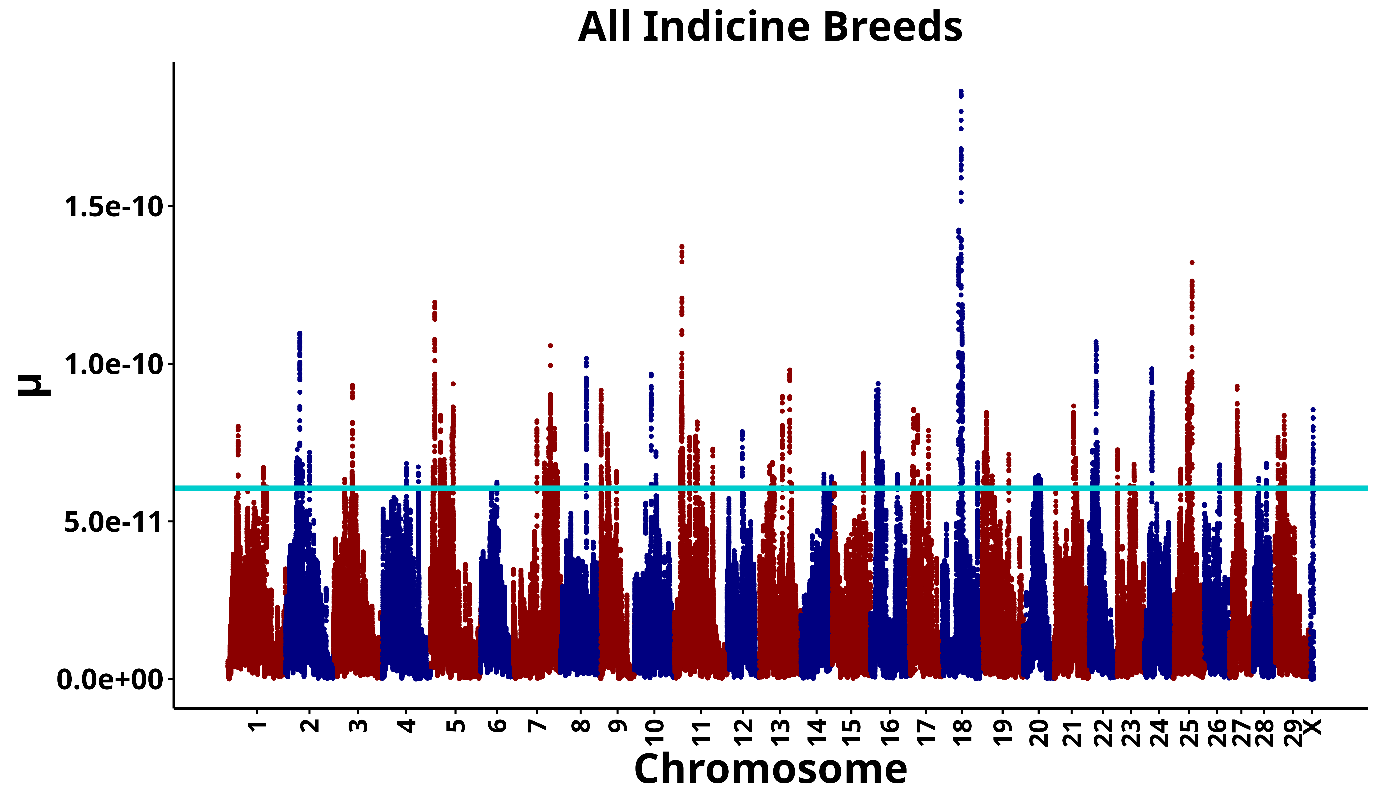
