## Supplementary Tables for "Integrative Whole-Genome Analysis reveals Genomic Signatures of Innate Immunity in Indicine Cattle"

**Supplementary Table 1: Sampling details for six cattle breeds across three different locations.** The table lists the breed, sampling location, and sample type used for DNA isolation and subsequent whole genome sequencing

| **Breed** | **Location** | **Sample** |
| --- | --- | --- |
| KangayamL1 | Thiruvaiyar | Blood |
| KangayamL2 | Vellaikovil, Kangayam | Blood |
| KangayamL3 | TANUVAS, Namakal | Blood |
| GirL1 | ICAR-NDRI, Karnal | Blood |
| GirL2 | Govindapuram, Kumbakonam | Blood |
| GirL3 | Bhopal | Blood |
| TharparkarL1 | ICAR-NDRI, Karnal | Blood |
| TharparkarL2 | ICAR-IVRI | Blood |
| TharparkarL3 | Babugarh | Blood |
| SahiwalL1 | ICAR-NDRI, Karnal | Blood |
| SahiwalL2 | Kalsi | Blood |
| SahiwalL3 | Babugarh | Blood |
| Karan FriesL1 | ICAR-NDRI, Karnal | Blood |
| Karan FriesL2 | - | Fresh semen |
| Karan FriesL3 | - | cryopreserved semen  (7 years and 12 years ago) |
| Holstein FriesianL1 | Lucknow | Blood |
| Holstein FriesianL2 | Hisar | Blood |
| Holstein FriesianL3 | Babugarh | Blood |

**Supplementary Table 2:** Statistical summary of the whole genome sequencing data. The table presents number of reads, Q20, and Q30 scores of raw and trimmed sequence reads, highlighting the quality is improved for the reads trimmed with Trimmomatic.

| **Samples** | **Orientation** | **Before Trimming (Raw data sets)** | | | **After Trimming (Using Trimmomatic)** | | |
| --- | --- | --- | --- | --- | --- | --- | --- |
|  |  | **# reads** | **Q20 %** | **Q30 %** | **# reads** | **Q20 %** | **Q30 %** |
| KangayamL1 | R1 | 70643378 | 95.58 | 89.47 | 68953922 | 98.3 | 93.4 |
| KangayamL1 | R2 | 70643378 | 96.95 | 91.9 | 68953922 | 98.8 | 94.6 |
| KangayamL2 | R1 | 95782745 | 95.74 | 89.8 | 93344283 | 98.4 | 93.6 |
| KangayamL2 | R2 | 95782745 | 96.58 | 91.19 | 93344283 | 98.6 | 94.1 |
| KangayamL3 | R1 | 102594729 | 95.66 | 89.72 | 100201425 | 98.4 | 93.5 |
| KangayamL3 | R2 | 102594729 | 96.7 | 91.36 | 100201425 | 98.6 | 94.1 |
| GirL1 | R1 | 86818218 | 95.69 | 89.78 | 84593814 | 98.4 | 93.6 |
| GirL1 | R2 | 86818218 | 96.42 | 90.83 | 84593814 | 98.5 | 93.7 |
| GirL2 | R1 | 81045704 | 95.58 | 89.55 | 79245218 | 98.3 | 93.5 |
| GirL2 | R2 | 81045704 | 97.15 | 92.29 | 79245218 | 98.8 | 94.8 |
| GirL3 | R1 | 67377662 | 95.47 | 89.73 | 65349318 | 98.4 | 93.9 |
| GirL3 | R2 | 67377662 | 95.8 | 90.05 | 65349318 | 98.4 | 93.7 |
| TharparkarL1 | R1 | 91331895 | 95.75 | 89.78 | 87838753 | 98.4 | 93.6 |
| TharparkarL1 | R2 | 91331895 | 94.34 | 87.11 | 87838753 | 97.7 | 91.7 |
| TharparkarL2 | R1 | 77763116 | 95.08 | 88.88 | 75793655 | 98.3 | 93.3 |
| TharparkarL2 | R2 | 77763116 | 96.54 | 91.06 | 75793655 | 98.6 | 94.1 |
| TharparkarL3 | R1 | 74005339 | 95.67 | 89.63 | 71222322 | 98.3 | 93.4 |
| TharparkarL3 | R2 | 74005339 | 94.22 | 86.91 | 71222322 | 97.7 | 91.6 |
| SahiwalL1 | R1 | 75760775 | 94.83 | 88.26 | 72548258 | 98.2 | 93.1 |
| SahiwalL1 | R2 | 75760775 | 94.99 | 88.25 | 72548258 | 97.9 | 92.2 |
| SahiwalL2 | R1 | 115686635 | 95.39 | 89.24 | 112504117 | 98.3 | 93.4 |
| SahiwalL2 | R2 | 115686635 | 96.65 | 91.51 | 112504117 | 98.7 | 94.6 |
| SahiwalL3 | R1 | 79373112 | 95.55 | 89.38 | 75867986 | 98.3 | 93.3 |
| SahiwalL3 | R2 | 79373112 | 93.93 | 86.57 | 75867986 | 97.7 | 91.6 |
| Karan FriesL1 | R1 | 95713832 | 95.78 | 89.8 | 92253084 | 98.4 | 93.5 |
| Karan FriesL1 | R2 | 95713832 | 94.73 | 87.84 | 92253084 | 97.9 | 92.2 |
| Karan FriesL2 | R1 | 90163286 | 94.67 | 88.13 | 86739807 | 98.2 | 93.0 |
| Karan FriesL2 | R2 | 90163286 | 95.56 | 89.36 | 86739807 | 98.2 | 93.0 |
| Karan FriesL3 | R1 | 70085095 | 95.95 | 90.1 | 68448917 | 98.4 | 93.6 |
| Karan FriesL3 | R2 | 70085095 | 96.45 | 90.85 | 68448917 | 98.5 | 93.7 |
| Holstein FriesianL1 | R1 | 101229799 | 95.64 | 89.66 | 98057619 | 98.4 | 93.5 |
| Holstein FriesianL1 | R2 | 101229799 | 95.48 | 89.15 | 98057619 | 98.2 | 92.9 |
| Holstein FriesianL2 | R1 | 82938114 | 95.77 | 89.71 | 80321239 | 98.3 | 93.4 |
| Holstein FriesianL2 | R2 | 82938114 | 95.55 | 89.22 | 80321239 | 98.1 | 92.8 |
| Holstein FriesianL3 | R1 | 70783380 | 95.86 | 89.87 | 68586312 | 98.3 | 93.4 |
| Holstein FriesianL3 | R2 | 70783380 | 95.46 | 89.07 | 68586312 | 98.1 | 92.7 |

**Supplementary Table 3:** Number of genes having variants with high, moderate, modifier, and low impact identified by DELLY and annotated by snpEff

| **Breed** | **High** | **Moderate** | **Modifier** | **Low** |
| --- | --- | --- | --- | --- |
| **KGL1** | Del: 1224, Dup: 28  Inv: 22, Bnd: 68 | Del: 9, Dup: 3844 Inv: 4633 | Del: 323  Dup: 545, Inv: 3 | Del: 18  Dup: 12, Inv: 1 |
| **KGL2** | Del: 2879, Dup: 16  Inv: 27, Bnd: 110 | Del: 12, Dup: 2263 Inv: 4487 | Del: 649  Dup: 361, Inv: 1 | Del: 22  Dup: 9 |
| **KGL3** | Del: 4355, Dup: 32  Inv: 19, Bnd: 113 | Del: 11, Dup: 3629 Inv: 5422 | Del: 890  Dup: 421 | Del: 28  Dup: 15 |
| **GRL1** | Del: 2924, Dup: 18  Inv: 23, Bnd: 109 | Del: 9, Dup: 1221 Inv: 3721 | Del: 513  Dup: 245, Inv: 6 | Del: 11  Dup: 5 |
| **GRL2** | Del: 2849, Dup: 18  Inv: 26, Bnd: 62 | Del: 6, Dup: 4843 Inv: 5924 | Del: 581  Dup: 640, Inv: 1 | Del: 19  Dup: 7 |
| **GRL3** | Del: 776, Dup: 8  Inv: 15, Bnd: 50 | Del: 6, Dup: 1451 Inv: 5272 | Del: 212  Dup: 218, Inv: 2 | Del: 17  Dup: 4 |
| **TPL1** | Del: 1573, Dup: 20  Inv: 25, Bnd: 104 | Del: 10, Dup: 2333 Inv: 4459 | Del: 446  Dup: 441 | Del: 25  Dup: 11 |
| **TPL2** | Del: 3814, Dup: 21  Inv: 28, Bnd: 70 | Del: 4, Dup: 2386 Inv: 4086 | Del: 724  Dup: 446, Inv: 5 | Del: 19  Dup: 12 |
| **TPL3** | Del: 1070, Dup: 10  Inv: 20, Bnd: 65 | Del: 5, Dup: 1592 Inv: 3151 | Del: 308  Dup: 230, Inv: 5 | Del: 14  Dup: 2 |
| **SWL1** | Del: 2963, Dup: 18  Inv: 17, Bnd: 95 | Del: 10, Dup: 2447 Inv: 7863 | Del: 487  Dup: 453, Inv: 1 | Del: 17  Dup: 7 |
| **SWL2** | Del: 1258, Dup: 22  Inv: 21, Bnd: 127 | Del: 13, Dup: 2754 Inv: 3008 | Del: 429  Dup: 372, Inv: 4 | Del: 27  Dup: 6 |
| **SWL3** | Del: 2030, Dup: 18  Inv: 23, Bnd: 78 | Del: 9, Dup: 4022 Inv: 1599 | Del: 450  Dup: 524, Inv: 2 | Del: 19  Dup: 8 |
| **KFL1** | Del: 185, Dup: 9  Inv: 21, Bnd: 52 | Del: 4, Dup: 64  Inv: 5143 | Del: 162  Dup: 100, Inv: 3 | Del: 14  Dup: 5 |
| **KFL2** | Del: 728, Dup: 7  Inv: 13, Bnd: 41 | Del: 4, Dup: 809  Inv: 2898 | Del: 271  Dup: 170, Inv: 4 | Del: 10  Dup: 4, Inv: 1 |
| **KFL3** | Del: 303, Dup: 13  Inv: 4, Bnd: 26 | Del: 4, Dup: 707  Inv: 422 | Del: 221  Dup: 162 | Del: 13  Dup: 8 |
| **HFL1** | Del: 156, Dup: 19  Inv: 16, Bnd: 54 | Del: 6, Dup: 1071 Inv: 249 | Del: 151  Dup: 178, Inv: 3 | Del: 14  Dup: 9 |
| **HFL3** | Del: 536, Dup: 16  Inv: 8, Bnd: 31 | Del: 1, Dup: 1592 Inv: 1660 | Del: 142  Dup: 270, Inv: 2 | Del: 10  Dup: 8 |

**Supplementary Table 4:** Number of genes having variants with high, moderate, modifier, and low impact identified by GATK and annotated by snpEff

| **Breed** | **High** | **Moderate** | **Modifier** | **Low** |
| --- | --- | --- | --- | --- |
| **KGL1** | SNV: 1302, Ins: 812, Del: 1291 | SNV: 12159, Ins: 364, Del: 438 | SNV: 36528, Ins: 34557, Del: 34961 | SNV: 1302, Ins: 812, Del: 1291 |
| **KGL2** | SNV: 1486, Ins: 868, Del: 1185 | SNV: 12865, Ins: 428, Del: 499 | SNV: 36554, Ins: 35020, Del: 35213 | SNV: 18416, Ins: 982, Del: 1077 |
| **KGL3** | SNV: 1596, Ins: 909, Del: 1300 | SNV: 13172, Ins: 457, Del: 530 | SNV: 36570, Ins: 35067, Del: 35291 | SNV: 18720, Ins: 1030, Del: 1131 |
| **GRL1** | SNV: 1517, Ins: 870, Del: 1154 | SNV: 12793, Ins: 418, Del: 490 | SNV: 36536, Ins: 34893, Del: 35139 | SNV: 18357, Ins: 883, Del: 1045 |
| **GRL2** | SNV: 1412, Ins: 841, Del: 1294 | SNV: 12298, Ins: 364, Del: 450 | SNV: 36554, Ins: 34793, Del: 35082 | SNV: 17696, Ins: 917, Del: 1054 |
| **GRL3** | SNV: 977, Ins: 584, Del: 700 | SNV: 9575, Ins: 389, Del: 444 | SNV: 36147, Ins: 31530, Del: 32261 | SNV: 14079, Ins: 337, Del: 405 |
| **TPL1** | SNV: 1481, Ins: 843, Del: 1082 | SNV: 12752, Ins: 393, Del: 475 | SNV: 36522, Ins: 34854, Del: 35058 | SNV: 18352, Ins: 933, Del: 1061 |
| **TPL2** | SNV: 1384, Ins: 882, Del: 1370 | SNV: 12335, Ins: 363, Del: 438 | SNV: 36537, Ins: 34728, Del: 35110 | SNV: 17582, Ins: 840, Del: 1059 |
| **TPL3** | SNV: 1321, Ins: 742, Del: 956 | SNV: 12218, Ins: 377, Del: 399 | SNV: 36472, Ins: 34525, Del: 34721 | SNV: 17682, Ins: 811, Del: 881 |
| **SWL1** | SNV: 1371, Ins: 816, Del: 986 | SNV: 12523, Ins: 401, Del: 438 | SNV: 36459, Ins: 34511, Del: 34605 | SNV: 17910, Ins: 796, Del: 852 |
| **SWL2** | SNV: 1585, Ins: 1039, Del: 1679 | SNV: 13259, Ins: 446, Del: 569 | SNV: 36557, Ins: 35134, Del: 35385 | SNV: 18784, Ins: 1068, Del: 1163 |
| **SWL3** | SNV: 1398, Ins: 820, Del: 998 | SNV: 12691, Ins: 413, Del: 457 | SNV: 36492, Ins: 34793, Del: 34912 | SNV: 18087, Ins: 907, Del: 900 |
| **KFL1** | SNV: 1084, Ins: 672, Del: 889 | SNV: 10923, Ins: 287, Del: 340 | SNV: 36375, Ins: 33742, Del: 33977 | SNV: 15781, Ins: 674, Del: 737 |
| **KFL2** | SNV: 1105, Ins: 642, Del: 858 | SNV: 11248, Ins: 284, Del: 342 | SNV: 36396, Ins: 33874, Del: 34010 | SNV: 16176, Ins: 682, Del: 692 |
| **KFL3** | SNV: 2116, Ins: 1151, Del: 1435 | SNV: 15287, Ins: 519, Del: 627 | SNV: 36506, Ins: 35632, Del: 35686 | SNV: 20545, Ins: 1287, Del: 1272 |
| **HFL1** | SNV: 855, Ins: 565, Del: 802 | SNV: 9829, Ins: 271, Del: 328 | SNV: 36310, Ins: 3311, Del: 33268 | SNV: 14107, Ins: 610, Del: 643 |
| **HFL3** | SNV: 663, Ins: 499, Del: 669 | SNV: 7974, Ins: 191, Del: 214 | SNV: 35892, Ins: 31179, Del: 31367 | SNV: 11560, Ins: 460, Del: 471 |
